# Bile Acid Restrained T Cell Activation Explains Cholestasis Aggravated Hepatitis B Virus Infection

**DOI:** 10.1101/2022.02.14.480376

**Authors:** Chujie Ding, Yu Hong, Yuan Che, Tianyu He, Yun Wang, Shule Zhang, Jiawei Wu, Wanfeng Xu, Jingyi Hou, Lijuan Cao, Haiping Hao

**Affiliations:** State Key Laboratory of Nature Medicines, Key Laboratory of Drug Metabolism and Pharmacokinetic, China Pharmaceutical University, Nanjing 210009, China

**Keywords:** Bile acid, SOCE, T cell activation, Hepatitis B Virus

## Abstract

Cholestasis is a common complication of Hepatitis B Virus (HBV) infection, characterized by increased intrahepatic and plasma bile acid levels. Cholestasis was found negatively associated with hepatitis outcome, however; the exact mechanism by which cholestasis impact on anti-viral immunity and impede HBV clearance remains elusive. Here, we found that cholestatic mice are featured with dysfunctional T cell response, and bile acids inhibit the activation and metabolic reprogramming of CD4^+^ T cells. Mechanistically, bile acids disrupt intracellular calcium homeostasis via inhibiting mitochondria calcium uptake and elevating cytoplasmic Ca^2+^ concentration of CD4^+^ T cells, leading to STIM1 and ORAIL1 decoupling and impaired store-operated Ca^2+^ entry which is essential for NFAT signaling and T cell activation. Moreover, in a transgenic mouse model of HBV infection, it was confirmed that cholestasis compromised T cells activation resulting in poor viral clearance. Collectively, our results suggest that bile acids play pivotal roles in anti-HBV infection via controlling T cells activation and metabolism, and that targeting regulation of bile acids may be a therapeutic strategy for host virus defense.

## Introduction

Cholestasis is a common risk factor that complicated in enterohepatic diseases, especially in liver diseases including fibrosis, cirrhosis and hepatitis (Allen et al, 2011; Postnikova et al, 2012; Zhong et al, 2003). Clinical studies revealed that cholestasis is negatively associated with hepatis outcomes in chronic hepatitis B virus infection, and intrahepatic cholestasis of pregnancy (ICP) patients were more susceptible to Hepatitis B Virus (HBV) infection (Jiang et al, 2020). Notably, bile acids are overall accumulated in patients with hepatitis virus infection (Sang et al, 2021), and mice infected with HBV showed an apparent enhance of cholesterol 7a-hydroxylase (CYP7A1), the rate-limiting enzyme promoting the bile acids de novo synthesis (Oehler et al, 2014). However, the intrinsic mechanism underlying how cholestasis may aggravate virus induced hepatic pathology remains largely elusive.

Prospective studies show that cholestasis aggravating pathological processes is mediated by accumulated bile acids, in view of that bile acid playing multiple roles in innate and adaptive immune responses. However, the immunomodulatory mechanism of bile acids is very complicated, due to the structural and biological activity diversity of bile acids. For instance, in innate immunity, bile acids may act upon receptors to play anti-inflammatory roles. Bile acids activation of TGR5-cAMP-PKA axis is effective in halting inflammatory responses in monocytes, via restricting NLRP3 inflammasome activation and IL-1β production (Guo *et al*, 2016). But accumulated bile acids in cholestasis condition alternatively act as danger-associated molecular pattern (DAMP) to exert pro-inflammatory effect. We had previously identified that bile acids, especially DCA and CDCA, in cholestasis or sepsis condition, activate both signal 1 and 2 of NLRP3 inflammasome in differentiated macrophages, but not in monocytes in a dose-dependent manner (Hao *et al*, 2017). Moreover, excessively accumulated bile acids are endogenous stimuli inducing immune collapse and tissue failure, by activating Apaf-1/caspase-4(−11, murine) pyroptosome-mediated pyroptosis both in matured macrophages and hepatocytes, which underlies pathological mechanism of cholestatic liver failure (Xu *et al*, 2021). In addition, pathologic concentrations of bile acids also induce tissue injury by aggravating recruitment of neutrophils and NETosis-mediated cell death, via inducing pro-inflammatory cytokines secretion (Gujral *et al*, 2003).

In addition to innate immunity, emerging studies have revealed modulatory effect of certain bile acid species on adaptive immunity. Oxidative or isomeric derivatives of bile acids are active in shaping adaptive immune responses, by modulating helper T cell (Th) and regulatory T cell (Treg) homeostasis. For example, Oxo-bile acids (-LCA/DCA) inhibit the differentiation of Th17 cells by binding to transcription factor retinoid related orphan receptor-γt (RORγt) and isoalloLCA increases the differentiation of Treg cells through the production of mitochondrial reactive oxygen species (mitoROS) and subsequently increased expression of FOXP3, interpreting the causal link between gut microbiota dysbiosis and host susceptibility to inflammatory colitis (Hang *et al*, 2019; Song *et al*, 2020). Bile acid homeostasis also play key roles in regulating tumor immunity; transformation balance between primary and secondary bile acids plays a vital role in modulating NKT cells mediated antitumor effect in HCC model, via regulating the expression of CXCL16 which is essential for NKT cells recruitment to microenvironment (Ma *et al*, 2018). Despite these advances in deciphering how bile acids modulate both innate and adaptive immunity, very less is known how bile acids regulate virus elicited immunological adaptations and their relative outcome in the clearance of virus.

Collective studies claimed that the adaptive immune response, rather than innate immune response, plays a crucial role in controlling both viral clearance and hepatopathy. During the acute phase of HBV infection, effective T-cell response is vital for viral clearance and is characterized by active and sustained CD4^+^ and CD8^+^ T-cell responses. Specifically, naïve T cells were activated and differentiated into effector T cells and exert anti-viral effect by releasing IFN-γ and IL-2 or direct targeting infected hepatocytes. Coincidently, T cells undergo metabolic reprograming characterized by reduced OXPHOS rate and enhanced glycolysis activity, which provides essential bioenergetic and biosynthetic nutrients for T cell functions (Pearce *et al*, 2013). Upon TCR engagement, store-operated calcium entry (SOCE) triggered Ca^2+^ signaling is necessary for T cell activation and metabolic reprogramming which induces Ca^2+^ entry into the cell through calcium-release-activated calcium (CRAC) channels (Feske, 2007) activating calcineurin and nuclear factor of activated T cells (NFAT) translocation (Hogan *et al*, 2003). NFAT2 is an important nuclear transcription factor that is able to promote the transcription of activation and glycolysis related gene, such as IL-2, Glut1 and HK2 (Macian, 2005). In contrast, chronic HBV infection often featured with sluggish adaptive immune responses characterized by ‘T-cell exhaustion’, leading to HBV escape from immune control (Sandalova *et al*, 2010). Similarly with tumor immunity, T cell exhaustion in chronic HBV infection is also defined by reduced T cell activity and weak response, sustained expression of immune checkpoint molecules (such as programmed cell death-1(PD-1), lymphocyte activation gene-3 (LAG-3) and cytotoxic T lymphocyte-associated antigen-4 and CD244), as well as transcriptional and metabolic states that distinct from that of functional T cells (Wykes & Lewin, 2018).

Despite the pathological processes of HBV infection and gene replication as well as host immune determinants governing infection outcome have been largely uncovered, the precise role of bile acids on adaptive immunity during hepatitis virus infection remains poorly characterized. In this study, we revealed that bile acids suppressed T cells activation and metabolic reprogramming induced by T-cell receptor (TCR) signaling. Bile acids, especially deoxycholic acid (DCA) and chenodeoxycholic acid (CDCA), restrained T cell activation and metabolic reprogramming by negatively regulating calcineurin -NFAT2 signaling. Mechanically, bile acid blocked mitochondrial Ca^2+^ uptake and thus elevating cytosolic Ca^2+^ concentration, hindering Orail1-STIM1 coupling and subsequent SOCE triggered Ca^2+^ influx which is essential for NFAT2 mediated T cell activation. Further, in a mouse model of T cell activation induced by staphylococcal enterotoxin B (SEB), we confirmed that cholestatic mice induced by bile duct ligation (BDL) showed compromised T cell activation. Moreover, in a mouse model of HBV infection induced by rAAV-HBV1.3 transfection, BDL mice showed an impeded capability of viral clearance. Collectively, these findings indicate that bile acids act as an immunosuppressor by restraining T cell activation, explaining why cholestasis aggravates HBV infection is prognose of poor outcome, hinting to that targeting bile acids may provide a therapeutic option for HBV infection with cholestasis.

## Results

### Cholestasis impaires T cell response by inhibiting T cell activation

Adaptive immune system is believed to play a major role in viral clearance, and cholestasis results in a defective anti-viral defense (Lang *et al*, 2018). However, the intrinsic mechanism of bile acid inhibiting T cell mediated anti-viral infection remains unclear. To identify the phenotype that bile acid affects adaptive immune system, we developed a bile duct ligation (BDL) mouse model to mimic the pathological condition of cholestasis, and screened a panel of T cell subsets. BDL mice were characterized with dramatic reduction of CD4^+^ and CD8^+^ T cells in both mesenteric lymph node (MLN) and liver **(Fig 1A-C; Gating strategies in Appendix Fig S1A)**. To investigate how cholestatic mice undergoing T cell impairment, we analyzed a panel of markers indicating T cell subsets, including activation, differentiation and exhaustion related antigens. Both CD4^+^ and CD8^+^ T cells isolated from MLN and liver of BDL mice were evidenced by remarkable reduction of proportion of CD69 and CD25 positive cells, which are indicative of activated T cells, and instead exhibited high levels of suppressive antigen CTLA-4 (**Fig 1D-L; Fig EV1A-F; gating strategies in Appendix Fig S1B**). Population of differentiated T cells in MLN and liver of BDL mice were largely comparable to sham group, including effector T cells indicated by CD44^+^CD62L^-^ and memory T cells indicated by CD44^+^CD62L^+^ (**Fig EV1G-N; gating strategies in Appendix Fig S1C**), suggesting that cholestasis might not influence T cell differentiation. In contrast to MLN and liver in enterohepatic system which are enriched with high levels of bile acids, neither activation nor suppressive markers of T cells isolated from spleen and PBMC were affected by BDL (**Fig EV2**). These results indicate that cholestasis can induce a loss and/or exhaustion of T-cell activation.

**Figure 1.**
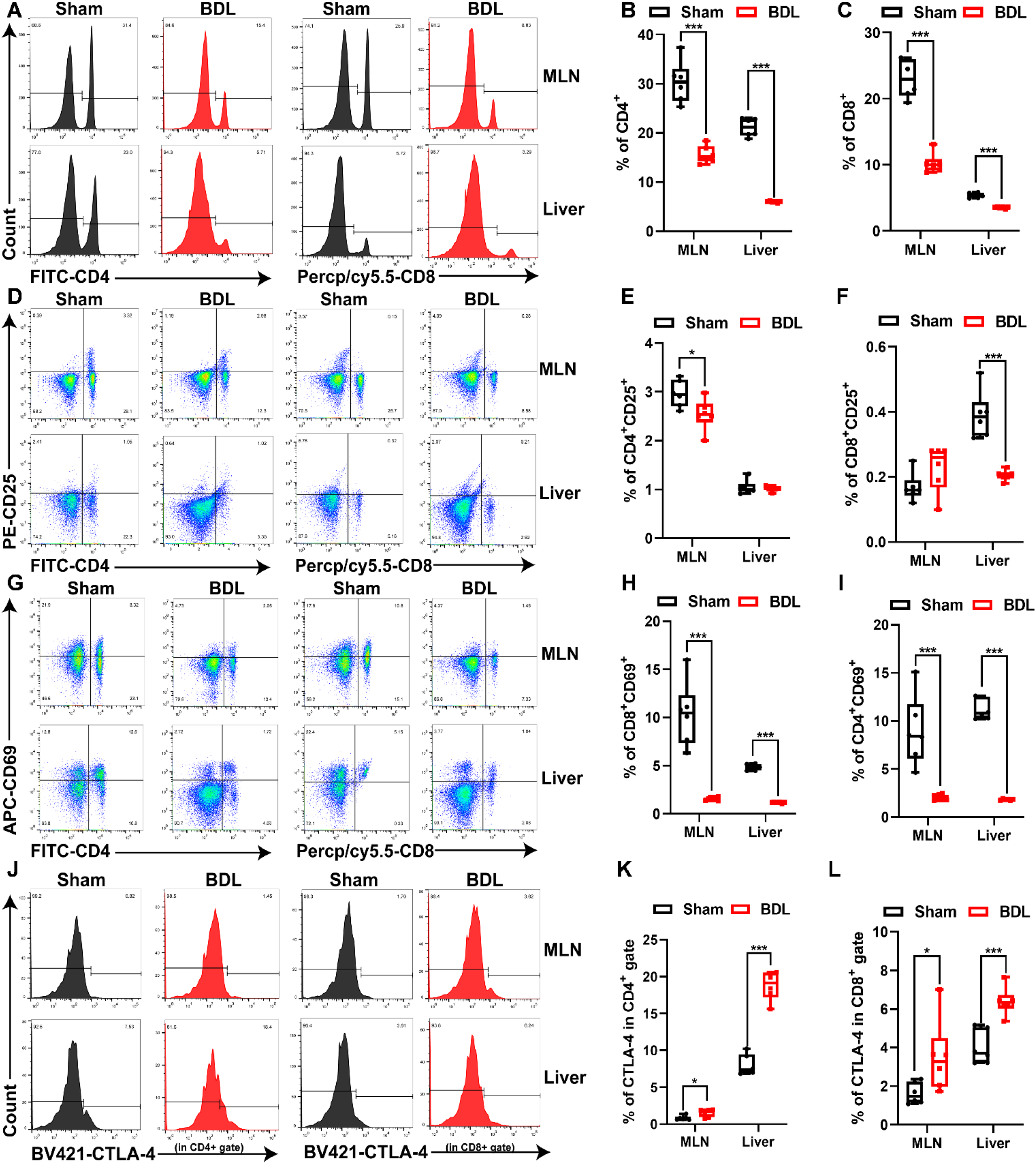
Cholestasis impairs T cell response by inhibiting T cell activation. Mice were performed with BDL surgery for 5 days before lymphocytes in MLN and liver were analyzed by flow cytometry. A-C Flow cytometry analysis (A) and quantitative data of CD4^+^ (B) and CD8^+^ (C) T cells in lymphocytes isolated from MLN and liver (*n* = 6). D-I Flow cytometry analysis (D, G) and quantitative data of activated CD4^+^ T cells (E, CD4^+^CD25^+^; H, CD4^+^CD69^+^) and activated CD8^+^ cells (F, CD8^+^CD25^+^; I, CD8^+^CD69^+^) in lymphocytes isolated from MLN and liver (*n* = 6). J-L Flow cytometry analysis (J) and quantitative data of CTLA among CD4^+^ (K) and CD8^+^ (L) T cells in lymphocytes isolated from MLN and liver (*n* = 6). Data information: Data are representative of three independent experiments and expressed as mean ± SEM. ***p<0.001, **p < 0.01, *p < 0.05 using two-tailed Student’s *t*-test. **See also Figure EV1 and EV2.**

To assess the pathological significance of reduced proportion of activated T cells under cholestatic condition, we next intraperitoneally injected with staphylococcal enterotoxin B (SEB) in mice to mimic pathogen induced T cell activation. Then the T cells activation were analyzed 6 hours after SEB injection. In comparison to sham control, BDL mice were insensitive to SEB induced T cell activation, as evidenced by decreased CD69^+^/CD25^+^ CD4^+^ and CD69^+^/CD25^+^ CD8^+^ populations in BDL mice (**Fig 2A-F**). Consistently, serum IL-2 and IFN-γ indicating T cell activation in response to SEB were much lower in BDL mice than that in sham mice (**Fig 2G and H**), demonstrating that cholestasis compromises both the CD4^+^ and CD8^+^ T cells activation induced by SEB. Taken together, these results indicate that cholestatic condition is unfavorable for T cell activation against pathogens in the acute stage of infection.

**Figure 2.**
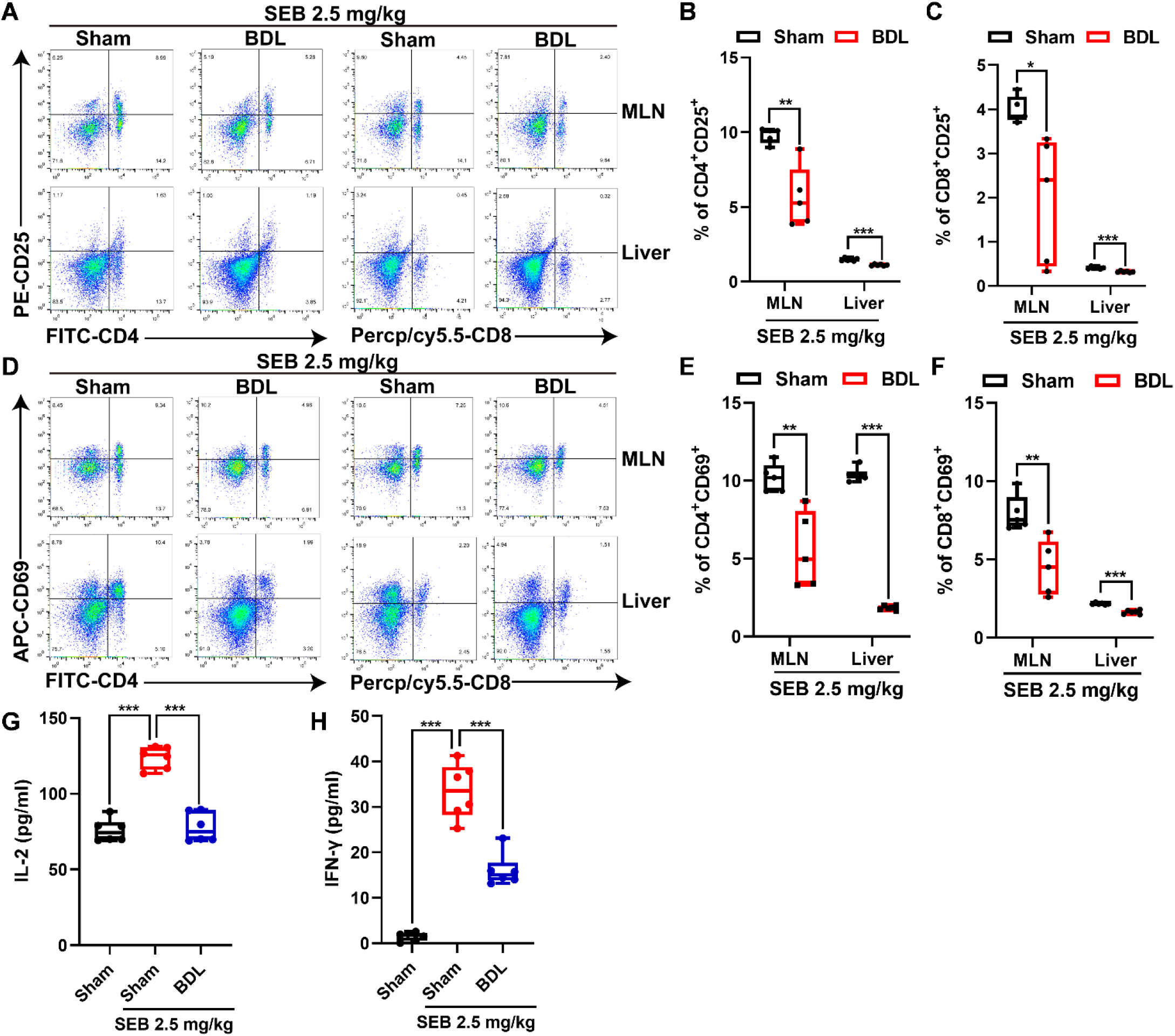
Cholestasis restrains SEB-induced T cells response *in vivo*. Mice were intraperitoneally injected with SEB (2.5 mg/kg mice body weight) at 5^th^ day after BDL surgery before MLN and liver were harvested 6 h later. A-F Flow cytometry analysis (A, D) and quantitative data of activated CD4^+^ T cells (B, CD4^+^CD25^+^; E, CD4^+^CD69^+^) and activated CD8^+^ cells (C, CD8^+^CD25^+^; F, CD8^+^CD69^+^) in lymphocytes isolated from MLN and liver (MLN, *n* = 5; liver, *n* = 6). G, H ELISA analysis of serum IL-2 (G) and IFN-γ (H) concentrations (*n* = 6). Data information: Data are representative of three independent experiments and expressed as mean ± SEM. ***p<0.001, **p < 0.01, *p < 0.05 using two-tailed Student’s *t*-test.

### Bile acids inhibit T cells activation by decoupling SOCE

We supposed that the elevated bile acid concentration might explain the observed effects of cholestasis in suppressing T cell activation. We screened a panel of bile acid species dominant in human and murine, by testing activation responses to anti-CD3/anti-CD28 antibodies stimuli with or without bile acid pretreatment in mouse primary CD4^+^ T cells. T cell activation was assessed by CD69 positive cell population. Notably CDCA, DCA, α-MCA, and their tauro-conjugates intensively suppressed T cell activation as evidenced by reduced CD69^+^ cell proportions (**Fig EV3A**). Moreover, DCA, CDCA, and α-MCA inhibited both early activation (indicated by CD69^+^ subsets) and late activation (indicated by CD25^+^ subsets) of mouse primary CD4^+^ T cell in a dose dependent manner, as well as subsequent IL-2 production (**Fig EV3B-L**), supporting bile acids are potent endogenous negative regulators of T cell activation. For further validation in human cells, Jurkat cells were stimulated with PMA/Ion with or without DCA/CDCA which are dominant species in human beings. As expected, DCA and CDCA significantly impaired Jurkat T cells activation, as shown by reduced CD69^+^ and CD25^+^ subset populations, as well as decreased IL-2 production in supernatant **(Fig 3A-D)**. IL-2 play a key role in driving T cell expansion, activation and differentiation to boost T cell response. Consistently, DCA and CDCA significantly inhibited the proliferation index of Jurkat cells, as assessed by CFSE labeling **(Fig 3E)**. Further, we validated regulatory role of DCA and CDCA on T cell activation in a mouse model of SEB-induced T cell activation. Both DCA and CDCA pretreatments significantly restrained CD4^+^ and CD8^+^ activation triggered by SEB, as evidenced by decreased CD69 and CD25 positive T cell subsets in MLN of mice. Consistently, IL-2 and IFN-γ was also decreased in bile acid pretreated mice (**Fig 3F-M; Fig EV4**). These results collectively demonstrate that bile acids, and especially DCA and CDCA, were active metabolites in controlling T cells activation both *in vitro and in vivo*.

**Figure 3.**
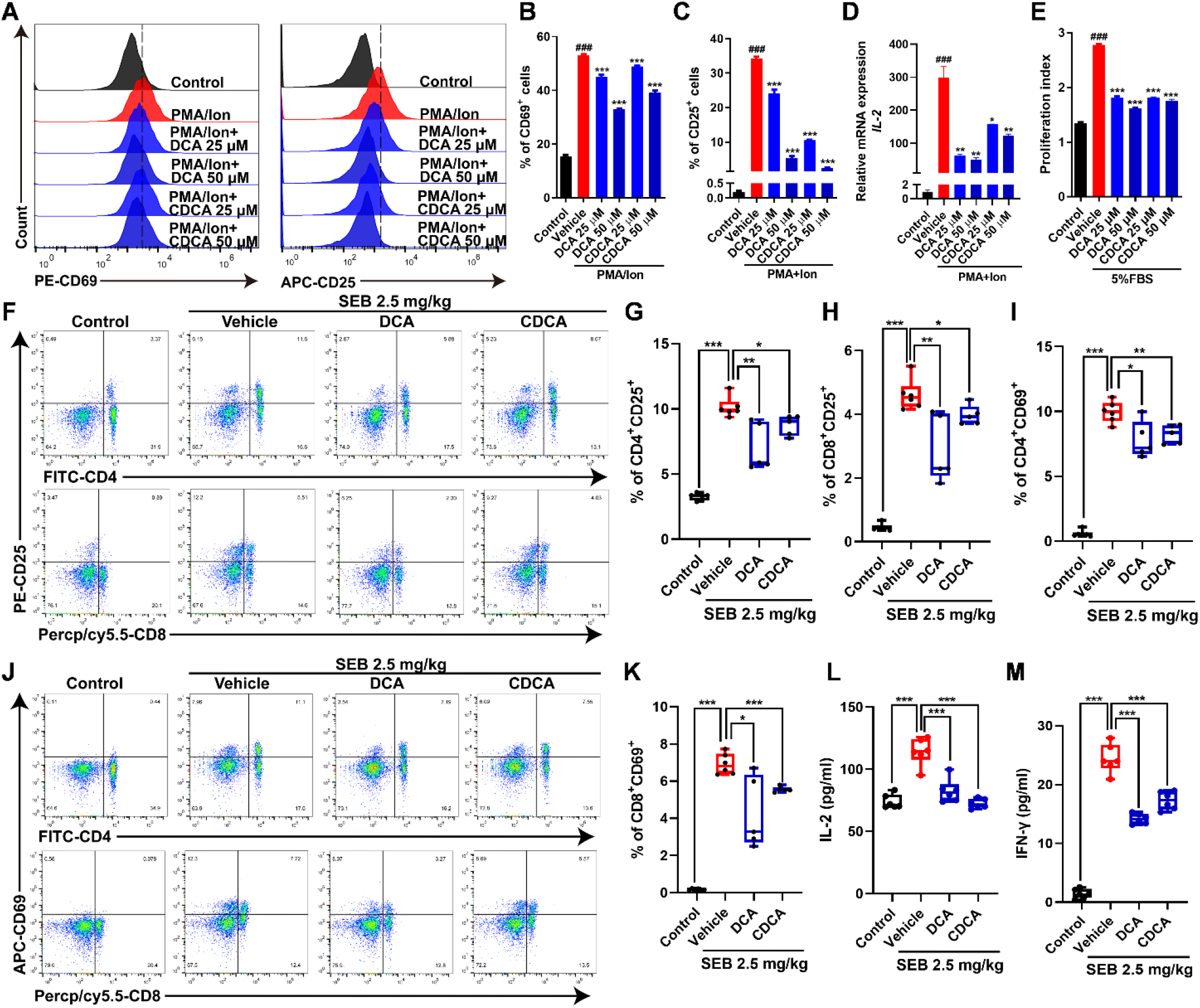
Bile acids inhibit T cells activation and proliferation. A-C Representative flow cytometry analysis and quantitative data of PMA/Ion stimulated Jurkat cells activation for 6 h (A, left; B) or 24 h (A, right; C) after bile acids pretreatment (25 or 50 µM, 24 h) (*n* = 3). D *IL-2* mRNA level detection by qRT-PCR in Jurkat cells which was pretreated with 25 and 50 µM BAs for 24 h and then stimulated with PMA/Ion for 6 h) (*n* = 3). E Proliferation index detected with CFSE in Jurkat cells treated with DCA and CDCA for 72 h (25 and 50 µM) (*n* = 3). F-K Representative flow cytometry analysis (F, J) and quantitative data of activated CD4^+^ T cells (G, CD4^+^CD25^+^; I, CD4^+^CD69^+^) and activated CD8^+^ cells (H, CD8^+^CD25^+^; K, CD8^+^CD69^+^) in lymphocytes isolated from MLN. Mice were pretreated with DCA or CDCA (25 mg/kg mice body weight, i.p.) for 7 days before SEB injection (2.5 mg/kg mice body weight, i.p.) for another 6 h (Control and Vehicle, *n* = 6; DCA and CDCA, *n* = 5). L, M ELISA analysis of serum IFN-γ (L) and IL-2 (M) concentrations in mice (*n* = 6). Data information: Data are representative of three independent experiments and expressed as mean ± SEM. (B-E) In comparison with the control group, ^###^p<0.001; compared with the vehicle group ***p<0.001, **p < 0.01, *p < 0.05 between the indicated groups; two-tailed Student’s *t*-test. (F-M) ***p<0.001, **p < 0.01, *p < 0.05 between the indicated groups using two-tailed Student’s *t*-test. **See also Figure EV3.**

We previously found in monocytes that bile acid modulates innate response via intervening intracellular calcium homeostasis (Hao et al., 2017). Thus we questioned if bile acids blunt T cells activation in a calcium signaling dependent manner. In contrast to monocytes which functional regulation is largely controlled by membrane channel mediated calcium influx, in T cells, SOCE represents the main source of calcium influx upon TCR signaling and is critical for calcineurin-NFAT activation. SOCE is mediated by Ca^2+^ release-activated Ca^2+^ (CRAC) channels that are activated by stromal interaction molecule (STIM) 1 and STIM2 which sense cytosolic calcium concentration alteration, and collective studies demonstrated that STIM1 and ORAI1 are the required components of SOCE (Wang *et al*, 2010). We thus intended to test whether bile acid could interfere SOCE signaling in T cells. Jurkat cells showed a remarkable interaction between STIM1 and ORAI1 upon PMA/Ion stimuli, while both DCA and CDCA significantly inhibited STIM1-ORAI1 coupling (**Fig 4A**). We then used thapsigargin (Tg) to deplete endoplasmic reticulum (ER) Ca^2+^ store and calcium chloride was added to trigger SOCE, and intracellular calcium was detected by use of Fluro-4 AM, consistently with STIM1-ORAI1 decoupling. Both DCA and CDCA pretreatment apparently abated calcium entry, supporting that bile acid was a negative regulator of SOCE signaling in T cells **(Fig 4B and C)**. In view that STIM1 activation and interaction with ORAI1 is triggered by calcium leak from ER and directly by elevated cytosolic Ca^2+^ we asked whether intracellular Ca^2+^ distribution was disturbed in bile acid preconditioned cells. Calcium leakage was barely changed in DCA/CDCA group in comparison with control group, as indicated by peak value of cytosolic calcium concentration ([Ca^2+^]_Cyto_) post Tg treatment (**Fig 4B**). We further analyzed subcellular Ca^2+^ distribution in Jurkat cells exposed to DCA and CDCA. Accompanied with elevated [Ca^2+^]_Cyto_, mitochondrial Ca^2+^ concentration ([Ca^2+^] _Mito_) was significantly decreased in bile acid treated cells (**Fig 4D and E**), indicating that bile acids disrupted calcium redistribution to the mitochondria in T cells upon calcium leakage from ER calcium pool. In T cells, mitochondria play a critical role in maintaining intracellular calcium homeostasis to reduce Ca^2+^-dependent slow inactivation (CDI) and decrease ER store refilling. The homeostasis between mitochondrial and cytosolic calcium is controlled by mitochondrial calcium influx and efflux pathways; influx is controlled by the activity of the mitochondrial Ca^2+^ uniporter (MCU), and efflux by the Na^+^/Ca^2+^exchanger (NCLX) and the H^+^/Ca^2+^ (mHCX) exchanger. (Hoth *et al*, 2000) (Baughman *et al*, 2011). We then assessed mitochondria Ca^2+^ uptake capacity in mitochondria isolated from bile acid treated Jurkat cells, mitochondria were challenged with successive additions of 50 mM CaCl_2_ and Ca^2+^ concentration were detected by Calcium Green-5N. In comparison to control group, mitochondria isolated from DCA and CDCA treated Jurkat T cells showed a significant decrease in calcium uptake rate and calcium retention capacity **(Fig 4F-I)**. Moreover, we verified MCU function by use of a MCU agonist spermine (Nicchitta & Williamson, 1984). Spermine pretreatment markedly halted inhibitory effects of bile acids on Tg/CaCl_2_ triggered SOCE **(Fig 4J-L)**, indicating that bile acids regulate intracellular Ca^2+^ redistribution in T cells mainly by inhibiting mitochondria Ca^2+^ uptake, and thereby leading to Ca^2+^ accumulation in cytosol and SOCE decoupling. To further confirm the impact of bile acids on SOCE and intracellular Ca^2+^ distribution *in vivo*, mice were intraperitoneally injected with DCA and CD4^+^ T cells were isolated from MLN for *ex vivo* assessment of SOCE. As expected, DCA led to an increase of [Ca^2+^]_Cyto_ and a decrease of [Ca^2+^]_Mito_ in CD4^+^ T cells from MLNs **(Fig 4M and N)**; consistently, the intensity of SOCE signaling was decreased in CD4^+^ T cells isolated from DCA-treated mice (**Fig 4O and P**). Taken together, these results indicate that mitochondrial calcium uptake controlled SOCE coupling might be a pivotal mechanism underlying inhibitory effect of bile acids on the T cell activation.

**Figure4.**
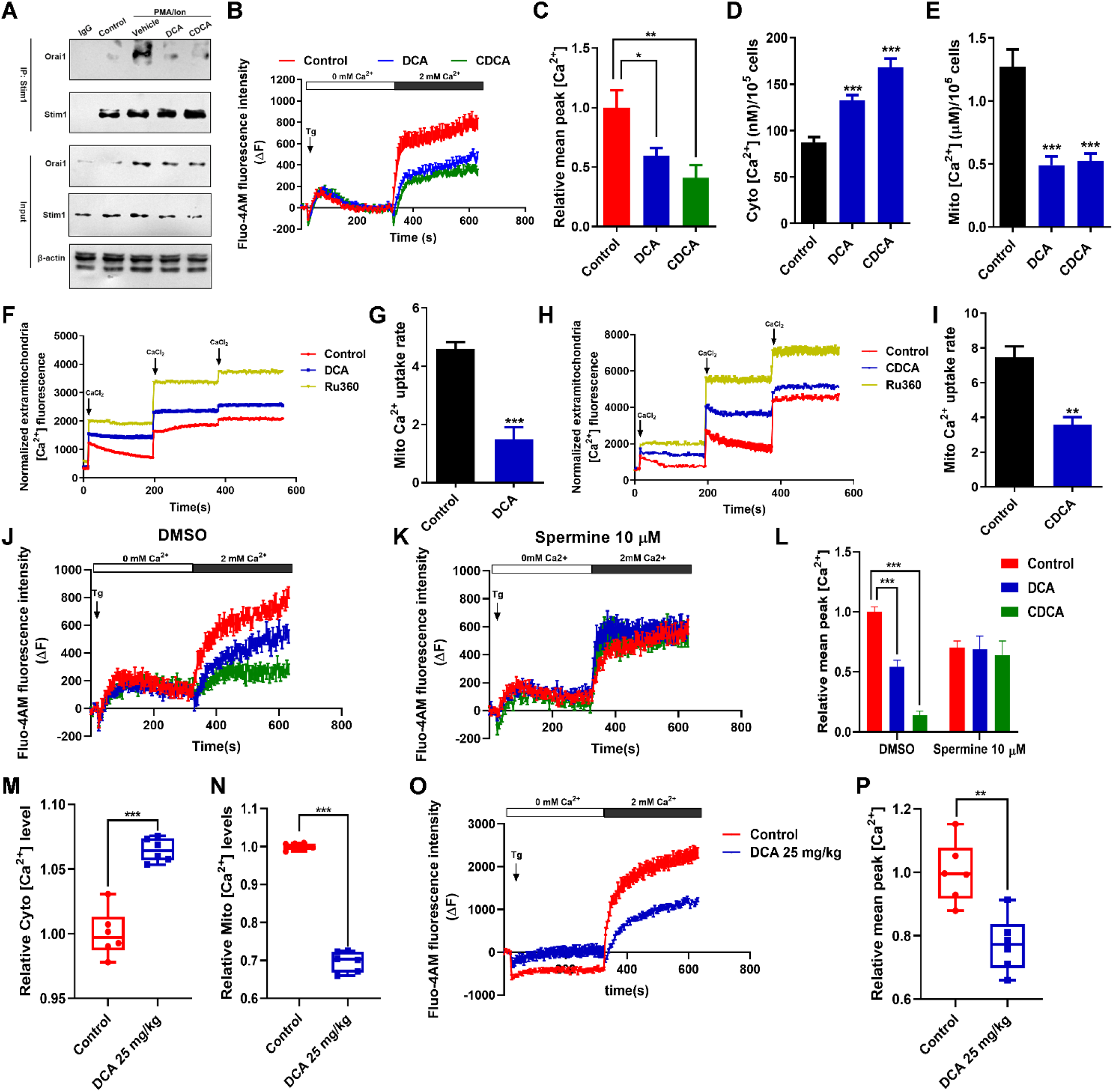
Bile acids suppress store-operated Ca^2+^ entry (SOCE) by reducing mitochondria Ca^2+^ exchange. A Co-IP analysis of the interaction between Stim1 and Orai1 in Jurkat cells lysates. Cells were pretreated with 25 µM DCA and CDCA for 24 h followed by stimulation with PMA/Ion for 1 h. B, C Jurkat cells were pretreated with 25 µM DCA and CDCA for 24 h. SOCE was detected using Fluo-4 AM upon 1 µM Tg stimulation without Ca^2+^ followed by re-addition of 2 mM extracellular Ca^2+^ (B) and quantitative data was shown in (C) (*n* = 6). D, E Cytosolic (D) and mitochondrial (E) Ca^2+^ concentration detection in Jurkat cells after DCA or CDCA treatment (25 µM, 24 h) using Fluo-4 AM and Rhod-2 AM (*n* = 6). F-I Mitochondria isolated from Jurkat cells were incubated with 25 µM DCA or CDCA for 30 min at room temperature. The effect of DCA or CDCA on the dynamics (F, DCA; H, CDCA) and rate (G, DCA; I, CDCA) of Ca^2+^ uptake in mitochondria from Jurkat cells was monitored using Calcium Green-5N. Ru360 was used as positive control. Each peak corresponds to 25 µM Calcium addition (*n* = 4). J-L Jurkat cells were exposed to 25 µM DCA or CDCA for 24 h and then preincubated with DMSO (J) or 10 µM Spermine (K) for 1 h before SOCE analysis using Fluo-4 AM. Quantitative data was shown in (L) (*n* = 6). M, N Relative cytosolic (M) and mitochondrial (N) Ca^2+^ levels of primary CD4^+^ T cells isolated from MLN in mice treated with DCA (25 mg/kg body weight) for 7 days (*n* = 6). O-P Mice were treated with DCA (25 mg/kg mice body weight) for 7 days. SOCE in primary CD4^+^ T cells isolated from MLN was detected using Fluo-4 AM and quantitative data was shown in (P) (*n* = 6). Data information: Data are representative of three independent experiments and expressed as mean ± SEM. ***p<0.001, **p < 0.01, *p < 0.05 using two-tailed Student’s *t*-test.

### Bile acids inhibit T cell activation and metabolic reprogramming depending on calcineurin-NFAT pathway

Upon TCR stimulation, SOCE evoked cytosolic Ca^2+^ elevation is crucial for calcineurin activation which dephosphorylates NFAT and promotes NFAT translocation to the nucleus to exert transcriptional activity (Vaeth & Feske, 2018). We therefore investigated the impact of bile acids on calcineurin-NFAT signaling pathway in T cells. In Jurkat T cells, PMA/Ion stimulation promoted phosphorylation of serine/threonine residues and phosphatase activity of calcineurin, which were inhibited by bile acids (**Fig 5A and B**). As the expression of NFAT2 is inducible and the expression of NFAT1 is constitutive, TCR signaling induces T cell activation mainly through induced transcriptional activity of NFAT2 (Macian, 2005). We then investigated NFAT2 translocation in Jurkat T cells with or without bile acid pretreatment. PMA/Ion stimulation induced NFAT2 translocation to the nucleus, while both DCA and CDCA strongly reduced the nucleus localization of NFAT2 as indicated by immunoblot of sub-cellular fractions **(Fig 5C)**. IL-2 is the major autocrine cytokine transcribed by NFAT and is necessary for T cells activation (Boyman & Sprent, 2012), accordingly, the level of IL-2 upon PMA/Ion stimulation was decreased in DCA and CDCA conditioned Jurkat T cells, in comparison to that of the control group **(Fig 3D)**.

**Figure5.**
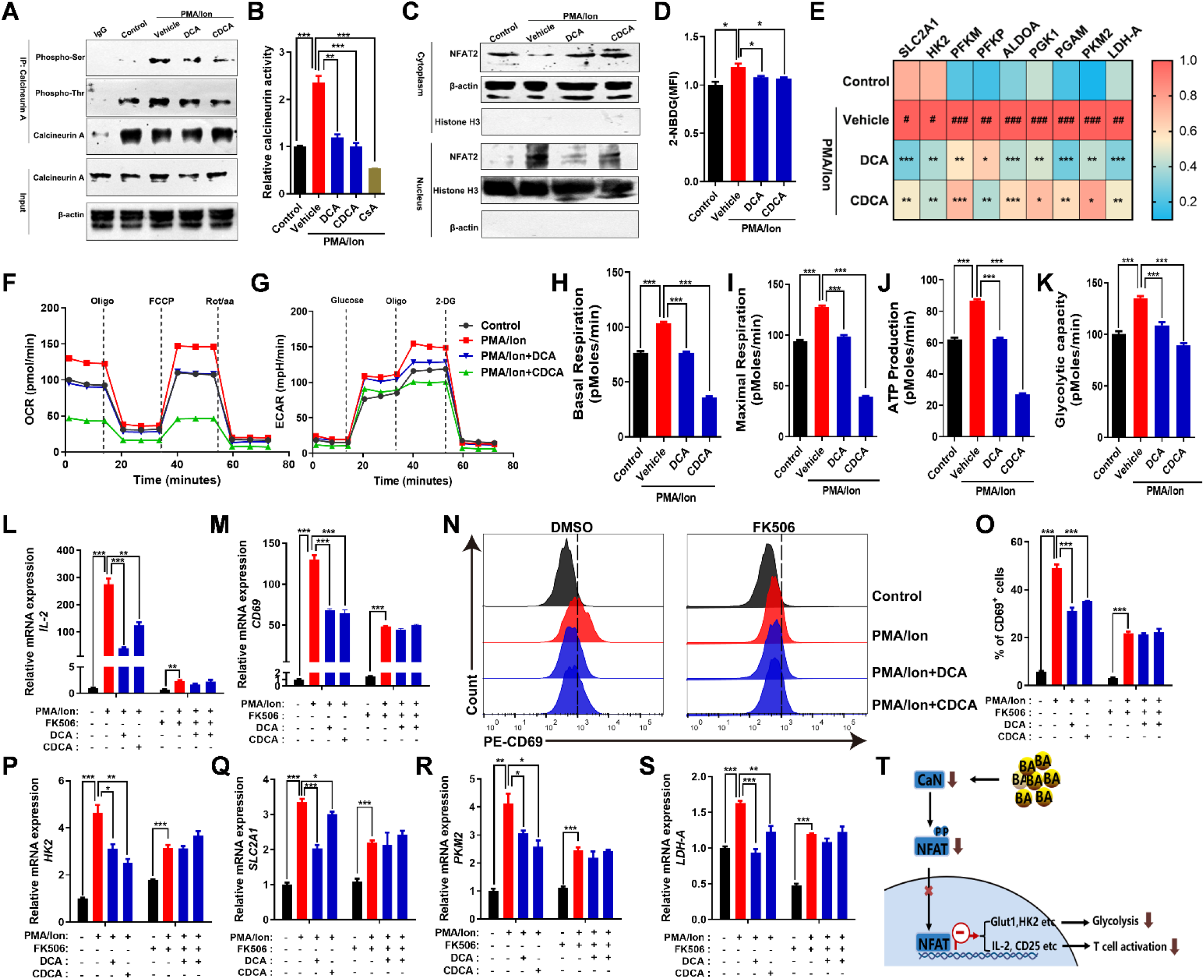
Bile acids regulate T cells activation and metabolism through calcineurin-NFAT pathway. A Phosphothreonine and phosphoserine detection of Calcineurin in Jurkat cells pretreated with DCA or CDCA (25 µM, 24 h) and PMA/Ion treatment for 3 h. B The calcineurin activity detection in Jurkat cells. Jurkat cells were pretreated with 25 µM DCA and CDCA for 24 h, then stimulated with PMA/ Ion for 1 h. 2 mM CsA was used as the positive control (*n* = 3). C The NFAT2 levels in the nuclear and cytosolic fractions were determined by immunoblotting. Jurkat cells were treated as described in (A). D Glucose uptake detection by 2-NBDG in Jurkat cells which was pretreated with DCA or CDCA (25 µM, 24 h) and stimulated with PMA/Ion for 1 h (*n* = 3). E Heatmap illustrating glycolytic enzymes mRNA changes in Jurkat cells. Jurkat cells were pretreated with DCA and CDCA 25 µM for 24 h and then stimulated with PMA and Ion for 6 h. F-K Extracellular flux analysis of oxygen consumption rate (OCR) (F), extracellular acidification rate (ECAR) (G), basal respiration (H), maximal respiration (I), ATP production (J) and glycolytic capacity (K) of Jurkat cells. Jurkat cells were treated as described in (E) (*n* = 8). L, M *IL-2* (L) and *CD69* (M) mRNA levels detection by qRT-PCR in Jurkat cells which was pretreated with 25 µM DCA or CDCA for 24 h and then stimulated with PMA/Ion for 6 h in the presence of 2 µM FK506 or DMSO (*n* = 3). N, O Representative flow cytometry analysis (N) and quantitative data (O) of CD69 in Jurkat cells which was pretreated with 25 µM DCA or CDCA for 24 h and then stimulated with PMA/Ion for 6 h in the presence of 2 µM FK506 or DMSO (*n* = 3). P-S The mRNA levels detection of glycolytic enzymes *HK2* (P), *SLC2A1* (Q), *PKM2* (R) and *LDH-A* (S) by qRT-PCR in Jurkat cells treated as described in (L, M) (*n* = 3). T Schematic illustration of the mechanism by which bile acids inhibit T cells activation and alter metabolism through calcineurin-NFAT pathway. Data information: Data are representative of three independent experiments and expressed as mean ± SEM. (A-D, F-S) ***p<0.001, **p < 0.01, *p < 0.05 between the indicated groups using two-tailed Student’s *t*-test. (E) In comparison with the control group, ###p<0.001, ##p < 0.01, #p < 0.05; compared with the vehicle group, ***p<0.001, **p < 0.01, *p < 0.05 between the indicated groups using two-tailed Student’s *t*-test. **See also Figure EV4.**

In addition to cytokines, NFAT also transcriptionally regulates glycolytic genes which is also a hallmark of T cell activation and is essential for T cells metabolic reprogramming to meet the supports for cell proliferation, activation and differentiation (Vaeth *et al*, 2017). We thus futher analyzed Jurkat cells metabolic activity by detecting glucose uptake activity with a glucose analog 2-NBDG and transcriptional profile of glycolytic genes. It was found that PMA/Ion boosted Jurkat cells metabolism while bile acids inhibited glucose metabolism, as indicated by the decrease of 2-NBDG uptake rate and the mRNA levels of glycolytic genes, including *SLC2A1*, hexokinase 2 (*HK2*), phosphofructokinase muscle (*PFKM*), pyruvate kinase isozymes M2 (*PKM2*), and *LDH-A, et al* (**Fig 5D and E**). Accordingly, we performed a non-targeted metabolomic study of Jurkat cells treated with DCA. DCA-conditioned cells were resistant to PMA/Ion induced metabolic reprogramming, particularly in TCA cycle, pyruvate metabolism, and pentose phosphate pathway (PPP) pathways **(Fig EV4A-E)**, suggesting a repression of glycolysis metabolism in bile acid treated T cells in response to PMA/Ion stimulation. To directly detect T cell metabolic reprogramming, we further assessed oxygen consumption (OCR) and extracellular acidification (ECAR) rates in Jurkat cells by seahorse analysis, indicative of OXPHOS and glycolysis, respectively. We found that PMA/Ion-activated Jurkat cells showed a drastic increase in OCR and ECAR rate and this increase was severely impaired by DCA and CDCA, indicated by basal respiration, maximal respiratory, ATP production, and glycolytic capacity **(Fig 5F-K)**. Consistent data were observed in CD3/CD28-activated primary mouse CD4^+^ T cells; energy metabolism of activated primary mouse CD4^+^ T cells largely relied on glycolysis represented by ECAR rate, which can be also abrogated by bile acids (**Fig EV4F-J**). Collectively, these results suggest that bile acids extensively inhibit T cell activation related metabolic reprogramming.

We next asked whether bile acids inhibited T cell activation and metabolic reprogramming were dependent of calcineurin-NFAT pathway. For further validation, we used an selective calcineurin inhibitor FK506 to prevent calcineurin-NFAT signaling pathway(Shaw *et al*, 1995),and we tested T cell activation response by detecting a panel of T cell activation and metabolic reprogramming markers, including relative mRNA levels of *IL-2, CD69, SLC2A1, HK2, PKM2* and *LDH-A* as well as percent of CD69^+^ positive cell populations. Bile acids impaired T cell activation response to PMA/Ion stimulation was abolished by FK506 pretreatment **(Fig 5L-S)**, supporting that SOCE-calcineurin-NFAT signaling pathway might be involved in bile acids repressed T cell activation and the associated metabolic reprogramming **(Fig 5T)**.

### Experimental cholestasis exacerbates HBV infection via repressing T cell activation

To further validate if bile acids-restrained T cell activation make sense in acute HBV infectious state *in vivo*, we next reproduced a mouse model of rAAV8-1.3HBV transfection to mimic acute phase of HBV infection. In line with clinical pathology, rAAV8-1.3HBV induced an instantaneous T cell activation response and liver injury within 1 day, followed by T cell exhaustion and the resultant increase of HBsAg concentration boosted at 7^th^ day **(Fig6A, Fig EV5A-F)**. We also observed that rAAV8-1.3HBV transfected mice were featured with a slight increase of serum total bile acid concentration at 7^th^ day. BDL mice were more susceptible to HBV infection as indicated by antigen clearance detected by serum HBsAg concentration and IHC of HBsAg in liver tissues, liver injury detected by HE staining, and serum AST and ALT levels (**Fig 6B-F**). Accordingly, BDL mice were characterized with attenuated activation of both CD4^+^ and CD8^+^ T cells triggered by rAAV8-1.3HBV, as indicated by decreased CD25 and CD69 positive CD4/8 subset populations, and reduced IFN-γ and IL-2 concentrations in the liver (**Fig EV5G-O, Fig 6G and H**).To interrogate whether the exacerbated HBV infection in BDL mice was mediated by T cells, we isolated CD8^+^ T cells from sham or BDL mice and co-cultured with Hepa1-6 cells and the co-culture system was simultaneously conditioned with rAAV-HBV1.3. We found that CD8^+^ T cells from sham mice were active in defensing against HBx invasion of hepatocytes; in contrast, CD8^+^ T cells from BDL mice showed compromised defense against HBx invasion of hepatocytes (**Fig 6I**). Furthermore, we depleted both CD4^+^ T cells and CD8^+^ T cells with anti-CD4 and anti-CD8 antibodies in sham and BDL mice prior to transfection with rAAV-HBV1.3 or rAAV-vector **(Fig 6J, Fig EV5O)**. Strikingly, mice with dual depletion of CD4^+^ and CD8^+^ T cells were more susceptible to HBV infection, but at this condition BDL induced cholestasis had negligible effects in HBV infection, as indicated by serum HBsAg levels and IHC of HBsAg in liver tissue at 14^th^ day of rAAV-HBV1.3 transfection (**Fig 6K and L**). Taken together, these data indicate that cholestasis aggravated HBV infection is related to the repression of T cell activation.

**Figure6.**
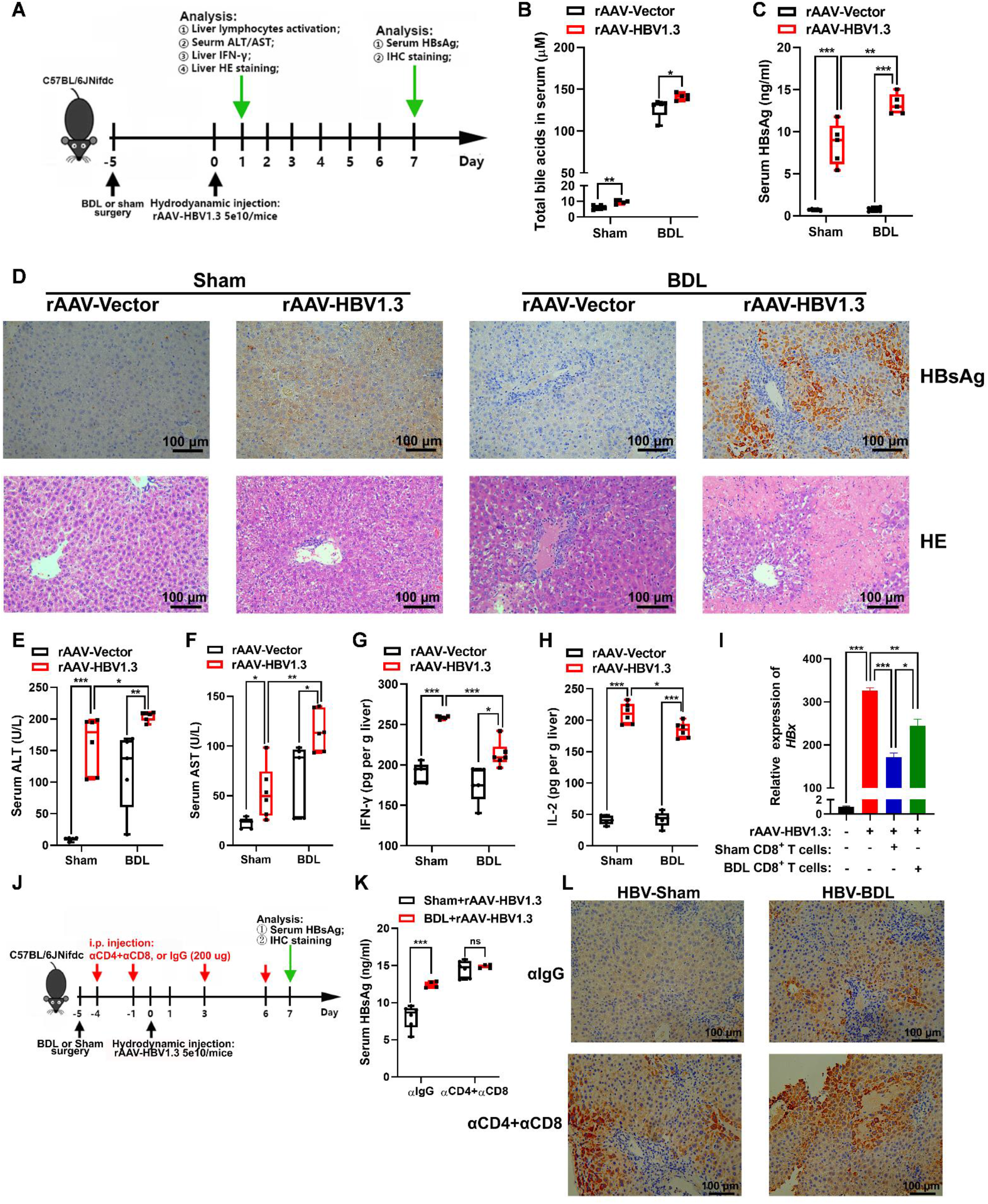
Cholestasis boosts HBV infection in a T cell-dependent manner. A Experimental scheme of BDL or sham mice injected with rAAV-HBV1.3 and rAAV-Vector. B Concentration of total bile acids in sham or BDL mice serum at 7^th^ day after rAAV-HBV1.3 virus transfection. (*n* = 6). C The serum HBsAg levels detection in sham or BDL mice at 7^th^ day after rAAV-HBV1.3 virus transfection (*n* = 5). D Representative HBsAg IHC staining from mice liver at 7^th^ day (upper) and H&E staining (bottom) from mice liver at 1^st^ day after rAAV-HBV1.3 transfection. Scale bar, 100 µm. E, F Mice serum ALT (E) and AST (F) levels detection at 1^st^ day after rAAV-HBV1.3 virus transfection following BDL surgery for 5 days (rAAV-Vector, *n* = 5; rAAV-HBV1.3, *n* = 6). G, H The IFN-γ (G) and IL-2 (H) concentrations detection in mice liver at 1^st^ day after rAAV-HBV1.3 virus transfection by ELISA (rAAV-Vector, *n* = 6; rAAV-HBV1.3, *n* = 5). I CD8^+^ T cells isolated from sham or BDL mice were co-cultured with Hepa1-6 cells and then transfected with rAAV-HBV1.3 for 48 h. The relative expression of *HBx* mRNA in Hepa1-6 cells was analyzed by qRT-PCR (*n* = 3). J Experimental scheme of BDL or sham mice injected with rAAV-HBV1.3 and rAAV-Vector with treatment of αCD4+αCD8 depleting antibodies in combination. K The serum HBsAg levels detection in mice as described in (J) (Sham+rAAV-HBV1.3, *n* = 6; BDL+rAAV-HBV1.3, *n* = 4). L Representative HBsAg IHC staining from mice liver as described in (J). Scale bar, 100 µm. Data information: Data are representative of three independent experiments and expressed as mean ± SEM. ***p<0.001, **p < 0.01, *p < 0.05 using two-tailed Student’s *t*-test. **See also Figure EV5.**

## Discussion

Although it was established that cholestasis is a common complication of HBV infection and cholestatic patients were more susceptible to hepatitis virus infection (Ferrari *et al*, 1990), the underlying mechanism of how cholestasis complicated in the pathological development of HBV infection remains largely unknown. The current study demonstrates that bile acid functions as an immunosuppressive metabolite in the adaptive immunity under cholesteric conditions. Mechanistically, bile acids disrupt intracellular calcium redistribution upon TCR signaling-triggered calcium leak from ER Ca^2+^ store by inhibiting mitochondria calcium uptake, leading to Ca^2+^ accumulation in the cytosol and SOCE decoupling. Moreover, bile acid induced SOCE decoupling is responsible for its inhibitory effect on calcineurin-NFAT pathway during T cell activation, and consequently repress the transcriptional activity of cytokines and metabolic reprogramming related genes and thereby preventing the activation of T cells. Therefore, T cells under cholesteric conditions are deficient in defending against HBV infection in acute phase leading to more aggravated hepatitis pathology (**Fig 7**).

**Figure7.**
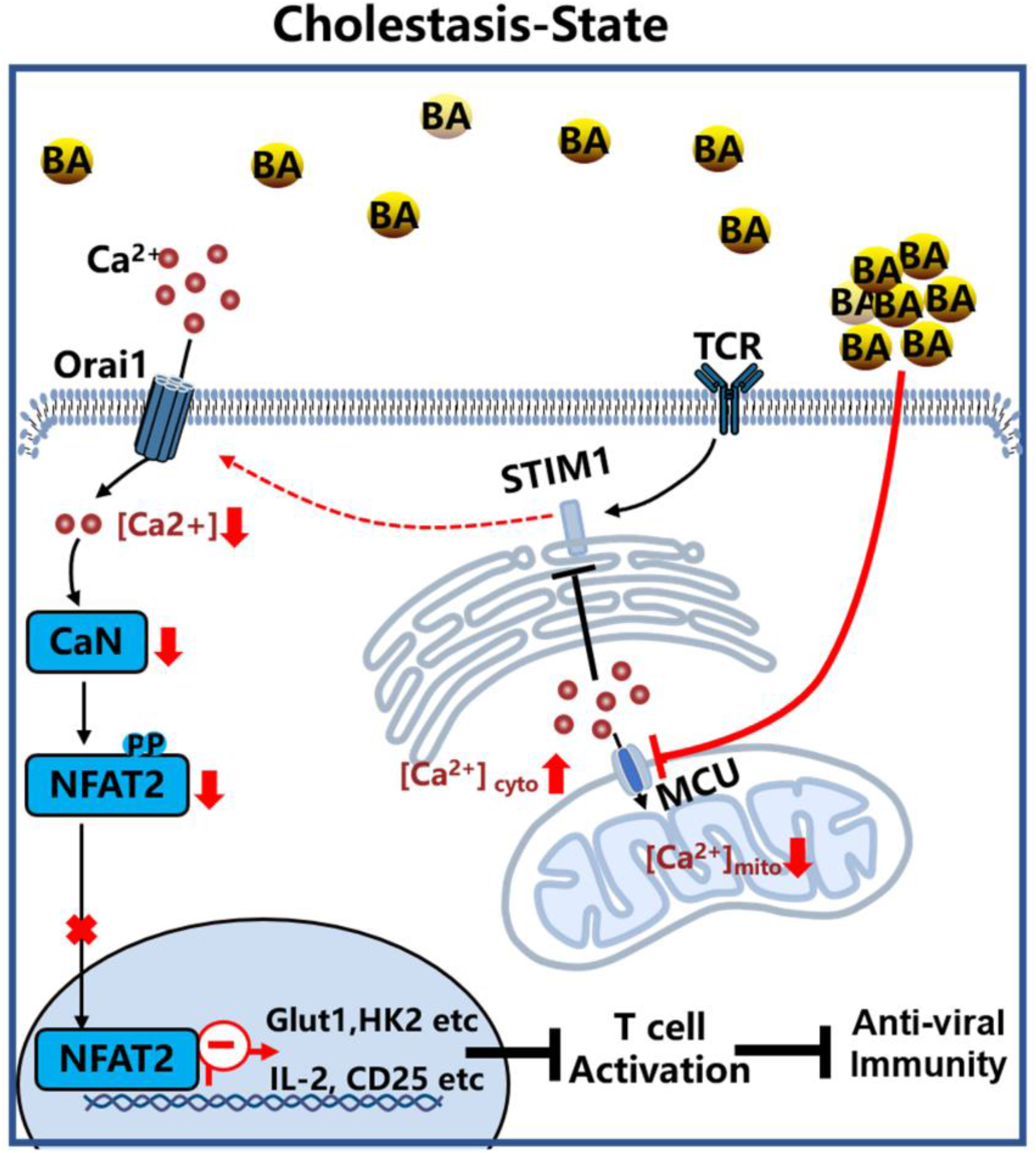
Diagram of bile acids impaired T cells activation to aggravate HBV infection.

Bile acids are derived from cholesterol in liver and further transformed by both host and gut microbiota. The imbalanced homeostasis of bile acids is recognized to be related to the occurrence of many diseases, including enterohepatic inflammation, infection, tumorigenesis, and diabetes (Haeusler *et al*, 2013; Hosseinkhani *et al*, 2021). We and others have disclosed that bile acids can be signaling molecules to exert diverse immunomodulatory activities on both innate immune and adaptive immune system, including pro-inflammatory effect on macrophages, activating pyroptosis in monocytes and NETosis in neutrophils, as well as modulatory effect on T cell differentiation (Hang et al., 2019; Hao et al., 2017; Xu et al., 2021). However, the detailed roles of bile acids on T cells during viral infection is not yet clear. A key observation of our study is that bile acids at pathologic level inhibited the activation of T cells, of note, DCA and CDCA as well as murine specific α-MCA are among the most active metabolites.

HBV infection caused hepatitis B is a serious disease affecting global public health (Terrault *et al*, 2016). However, the diagnosis and treatment of hepatitis B is still one of the most difficult public health problems at present. In clinical, nucleotide analogues and immunomodulators, such as interferon-α (IFN-α) are effective antiviral drugs approved by the FDA for the treatment of hepatitis B patients. Nucleotide analogues are reverse transcriptase inhibitors that can suppress HBV replication effectively, and thus are prescribed as the first line of treatment. But the long-term use the nucleotide analogues drugs may have drug resistance and side effects, which is the major challenge currently for HBV therapy (Guo *et al*, 2018). And once people start nucleotide analogues drugs, they must continue indefinitely. By contrast, IFN-α serves an antiviral cytokine and enhances the immune function of T cells to kill cells infected by virus. IFN-α has the advantages of shorter duration of administration and avoiding drug resistance; however, the current clinical application of IFN-α inhibits virus replication in hepatitis B patients only 20-40% (Hoofnagle *et al*, 2007). Existing hepatitis B treatment strategies cannot completely eliminate the HBV; therefore, it is necessary to propose more effective treatment options. Cohort studies found there was a strong positive correlation between hepatitis virus and ICP patients and previous HBV infection may aggravated the primary biliary cholangitis (PBC) severity, leading to poorly outcomes (Marschall *et al*, 2013; Zhang *et al*, 2018).Our study establishes that cholestasis impairs T cell immune activation to aggravate HBV infection and liver injury. This phenotype is akin to the increased susceptibility to HBV infection seen in cholestasis patients. Results obtained from our study suggest that targeting bile acids, for patients with HBV infection and the concurrent cholestasis, might be a promising strategy by restoring the response and activities of T cells in clearing infected virus. Notably, cholestasis is also a high incident complication for other virus infection related diseases. For instance, patients infected with COVID-19 may also progress to liver injury and cholestasis, and accompanied repressed T cell activation response. Findings of our study may ignite future research to explore whether bile acids repressed T cell activation response can be also involved in the pathological development of other virus infection related diseases.

## Materials and Methods

### Reagents and antibodies

PMA (P1585), Deoxycholic acid (D2510) and Chenodeoxycholic acid (C9377) were purchased from Sigma-Aldrich (St. Louis, MO, USA). Ionomycin (I139530) was purchased from Aladdin (Shanghai, China). FK506 (HY-13756) was purchased from MedChemexpress (Monmouth Junction, NJ, USA). LEAF Purified anti-mouse CD3ε Antibody (100313) and LEAF Purified anti-mouse CD28 Antibody (102111) were purchased from Biolegend (San Diego, CA, USA). Staphylococcus Entrotoxin B (BT202) was purchased from Toxin Technology (Sarasota, Fla). Viral genome rAAV8/HBV1.3 (serotype ayw) was purchased from Genewiz (Suzhou, China). InVivoMab anti-mouse CD4 (BE0003-1), InVivoMab anti-mouse CD8 (BE0061) and InVivoMab rat IgG2b isotype control (BE0090) were purchased from Bioxcell (West Lebanon, NH).

Antibodies used for western blot were from as the following, Calcineurin A (2614S), Histone H3 (4499S) and Stim1 (5668S) were purchased from Cell Signaling Technology (Danvers, MA, USA). NFAT2 (ab25916), Orai1 (ab111960), Phosphoserine (ab9332) and Phosphothreonine (ab9337) were purchased from Abcam (Cambridge, UK). β-actin (HC201-02) was purchased from Transgen biotech (Beijing, China). PP2B-Aα (sc-17808) and Stim1 (sc-66173) were purchased from Santa Cruz Biotechnology (Santa Cruz, CA, USA). Mouse IgG_1_(66360) was purchased from Proteintech (Chicago, USA). Mouse IgG_2b_ (400301) were purchased from Biolegend (San Diego, CA, USA).

For flow cytometry analysis, anti-mouse CD16/32(101302), FITC anti-mouse CD4 (100406), PerCP/cy5.5 anti-mouse CD8a (100734), PE anti-mouse CD25 (102007), APC anti-mouse CD69 (104514), PE anti-mouse IFN-γ (505807), APC anti-mouse CD25 (102001), Brefeldin A solution (420601) and CFSE Cell Division Tracker Kit (423801) were purchased from Biolegend (San Diego, CA, USA). PE anti-mouse IL-4 (12-7041-81), APC anti-mouse/rat IL-17A (17-7177-81), PE anti-mouse/rat Foxp3 (12-5773-82), PE anti-human CD69 (12-0699-42), APC anti-human CD25 (17-0259-42) and 2-NBDG (N13195) were purchased from Thermo Fisher Scientific (Waltham, MA, USA).

### Cell lines and culture

Human acute T cell leukemia cell line Jurkat T cell was obtained from Stem Cell Bank of the Chinese Academy of Sciences (Shanghai, China) and cultured in RPMI-1640 medium (Gibco, Pittsburgh, PA, USA) containing 10% Fetal Bovine Serum (FBS) and 100U/ml penicillin-streptomycin. Hepa1-6 cell line was obtained from Stem Cell Bank of the Chinese Academy of Sciences (Shanghai, China) and cultured in DMEM medium (Gibco, Pittsburgh, PA, USA) containing 10% Fetal Bovine Serum (FBS) and 100U/ml penicillin-streptomycin. All cells were grown at 37 °C in a humidified atmosphere containing with 5 % CO_2_. All cell culture medium were decontaminated from mycoplasma by following the instructions of Mycoplasma Detection Kit (Yeasen, Shanghai, China).

### Mice

Male C57BL/6JNifdc mice (6-7 weeks of age), obtained from Vital River Laboratories (Beijing, China). Mice were maintained under standard laboratory conditions of a constant temperature of 22 °C, a 12-h light-dark cycle, and given free access of drinking water and diet. The mice were group housed and acclimated for 7 days before the experiments began. All the animal experiments were approved by the Animal Ethics Committee of China Pharmaceutical University, and the procedures were performed in accordance with the Institutional Animal Research Committee guidelines to ensure minimal discomfort in the animals.

### Animal treatments

To investigate the role of DCA and CDCA on T cell activation in vivo, we through Staphylococcus Enterotoxin B (2.5 mg/kg, i.p.) to induce mouse T cell activation model, which commonly triggers T cell activation at∼6 h. For bile acids treatment, mice were treated with DCA and CDCA (25 mg/kg, i.p.) or sterile saline daily for 7 days before SEB injection. The spleen and mesenteric lymph nodes were harvested for further analyses.

To examine the impact of cholestasis on T cell activation in vivo, we had a bile duct ligation. All animals were anesthetized with chloral hydrate solution (0.3 g/kg) administered by intraperitoneal injection. After induction of anesthesia, a median abdominal incision was made and the common bile duct was identified. The duct was dissected carefully and doubly ligated with 7–0 Prolene (Ethicon, Somerville, NJ). In the sham operation group, the duct was dissected without ligation. The abdominal incision was closed in two-layer sand wiped with alcohol swab after surgery; the mice were kept warm until recovery. After five days of surgery, mice were injected intraperitoneally with Staphylococcus Enterotoxin B (2.5 mg/kg, i.p.) to induce mouse T cell activation model. The mesenteric lymph nodes were harvested for further analyses.

To test the effect of HFD on T cell activation in vivo, the mice were randomized into the HFD group (received 60% kcal fat diet) and CD group (received 10% kcal fat diet) at six weeks of age. After 12 weeks of treatment, mice were sacrificed, and the spleens and mesenteric lymph nodes were collected and analyzed by flow cytometry.

To establish acute HBV infected mice, 5×10^10^ viral genome rAAV-HBV1.3 was injected into the tail vain of each mouse in a volume of sterile saline that was equivalent to 10% of the body mass of the mouse within 5-8 s(Yang *et al*, 2002). Control animals received an isovolume injection of sterile saline. In other series of studies examining the potential effects of cholestasis on HBV viral clearance. After five days of bile duct ligation surgery, mice were hydrodynamic injected with 5×10^10^ viral genome rAAV-HBV1.3. Separate groups of mice were sacrificed on day 1 and day 7 after rAAV-HBV1.3 injection. Peripheral blood and livers were harvested for further analyses.

To depletion of T cells, mice were treated with 200 µg of InVivoMab anti-mouse CD8, InVivoMab anti-mouse CD4 or InVivoMab rat IgG_2b_ in 200 µl PBS intraperitoneally (i.p.) at twice a week. T cells depletion was checked by flow cytometry.

### Spleen and mesenteric lymph node lymphocytes isolation

In brief, mice were euthanized by rapid cervical dislocation and sterilized with 75% alcohol for 5 min. Placed lymph nodes and approximate 2 cm^3^ section of spleen onto a sterile dish, using 2.5 ml Syringe plug gently grinded the lymph nodes and spleens and filtered by 4 ml PBS (containing 2% FBS) through a 40-µm cell strainer. Collected and centrifuged cell suspension 600 g for 15 min at 4 °C. After centrifugation, the spleen cell mass was suspended into 1ml Red Blood Cell Lysis Buffer on the ice for 5 min. Centrifuged cell suspension 600 g for 5 min at 4 °C. After centrifugation, spleen and lymph nodes cell mass were suspended in PBS (containing 2% FBS), flow cytometry and seahorse were performed.

### Intrahepatic leukocytes isolation

In brief, mice were euthanized by rapid cervical dislocation and sterilized with 75% alcohol for 5 min. The murine liver was removed and pressed through 70-µm cell strainer. Collected and centrifuged cell suspension 50 g for 5 min at 4 °C. After centrifugation, the supernatant is the suspension of liver non-parenchymal cells. Then centrifuged liver non-parenchymal cells suspension 2000 rpm for 10 min at 4 °C. After centrifugation, the cells mass was resuspended in 3 ml 40% Percoll solution (17-0891-09; GE, Sweden), gently overlaid onto 2 ml 70% Percoll solution, and centrifuged at 1260 g for 30 min at 4 °C. The Intrahepatic leukocytes at the interface between Percoll solution. Collected cells and washed by PBS.

### Primary mouse CD4^+^ T cells Purification

CD4^+^ T cells were isolated from spleen and mesenteric lymph nodes single cells suspensions by positive separation using a Dynabeads FlowComp Mouse CD4 Kit (11461D, Thermo Fisher Scientific) according to manufacturer’s instructions. In vitro culture, CD4^+^ T cells were stimulated in RPMI 1640 medium (supplemented with 100 U/ml penicillin-streptomycin) with 10 µg/ml plate-bound anti-CD3 (clone 145-2C11), plus with 2 µg/ml anti-CD28 Abs (clone 37.51) in the presence of 25 µM DCA and CDCA.

### Flow cytometry

Immune cell from mouse lymphoid organs and liver were blocked with anti-CD16/32 antibodies (clone 93) for 10 min on the ice before staining the surface markers. Staining of surface markers with fluorescently labeled antibodies was performed at room temperature for 20 min in the dark. Samples were acquired on a Celesta flow cytometer using FACSDiva software (BD Biosciences, San Jose, CA) and analyzed with FlowJo software V10.

### Seahorse analysis

Oxygen consumption rates (OCR) and extracellular acidification rates (ECAR) were measured using an XFe96 Extracellular Flux Analyzer (Agilent, San Diego, CA, USA). Jurkat T cells were stimulated with 25 ng/ml PMA and 0.5 µg/ml Ionomycin for 6h in the pretreated with 25µM DCA and CDCA for 24 h, seeded at 1×10^5^ cells/well. Isolated CD4+ T cells from spleen and lymph nodes were activated with plate-bound anti-CD3 plus with soluble anti-CD28 Abs in the presence of 25µM DCA and CDCA for 24 h, seeded at 4×10^5^ cells/well. Before experiments, cells were resuspended in XF medium (Agilent) supplemented with 1 mM sodium pyruvate (Sigma-Aldrich), 10mM sodium glucose (Sigma-Aldrich) and 1 mM Glutamine (Sigma-Aldrich). OCR was monitored under basal conditions and followed treat with oligomycin (1.0 µM), FCCP (0.75 µM), rotenone (100 nM) and antimycin A (1 µM). ECAR was monitored followed treat with glucose (10 µM), oligomycin (1.0 µM) and 2-DG (50 µM). All cells were measured using the Seahorse XF Glycolysis Stress Test Kit (Agilent) and Seahorse XF Cell Mito Stress Test Kit (Agilent) according to manufacturer’s instructions.

### Glucose Uptake Measurement

To measure glucose uptake, Jurkat T were pretreated with 25 µM DCA and CDCA for 24 h, followed by incubated with glucose-free RPMI 1640 medium (Gibco) supplemented with 25 ng/ml PMA, 0.5 µg/ml Ion and 100 µM 2-NBDG for 1 h at 37 °C. The fluorescence of 2-NBDG taken up by Jurkat T cells were assessed by flow cytometry.

### CFSE dilution assay

Jurkat T cells were incubated in RPMI 1640 medium with 5% FBS in the presence of DCA and CDCA 25 µM containing with 5 µM CFSE for 72 h. CFSE dilution level was assessed by flow cytometry, the proliferation index (PI) was analyzed by Modfit LT.

### Metabolite Profiling

LC/MS analyses were conducted on an AB SCIEX Triple TOF™ 5600 mass spectrometer system (AB Sciex), which was coupled to a LC-30A Shimadzu LC system (Shimadzu). An XBridge BEH Amide HPLC column (100 mm× 4.6 mm, 3.5 µm) (Waters, Milford, MA) was used for compound separation at 40 °C. The mobile phase consisted of buffer A (95% 5 mM ammonium acetate buffer, pH adjusted to 9.5% acetonitrile) and buffer B (acetonitrile). The chromatographic gradient was run at a flow rate of 0.4 ml/min as follows: 0-3 min, 85% B; 3-6 min, 85-30% B; 6-15 min, 30-2% B; 15-18 min, 2% B; 18-19 min, 2-85% B; and 19-26 min, 85% B. MS data was acquired in negative ESI modes at a range of *m/z* 50 to 1000. Relative quantitation of polar metabolites was performed with MultiQuant software (AB SCIEX).

Jurkat T Cells and serum samples preparation for metabolomics analysis was performed as previously described(Wu *et al*, 2017). Jurkat T cells and serum were permeabilized with ice-cold methanol containing with 1.5 µg/ml 4-Chloro-Phenylalanine (Sigma-Aldrich) as internal standard (IS). After vibration and centrifugation at 18000 g for 10 min at 4 °C. The supernatant was collected and evaporated to dryness at room temperature under vacuum. The samples were resuspended in 100 µl of water prior to metabolomics analysis performed by HPLC-Triple-TOF/MS instrument (AB Sciex), and 20 µl was used for analysis.

### Measurements of intracellular and mitochondria Calcium concentrations

Intracellular calcium fluorescence was measured by 1 µM Fluo-4AM (Thermo Fisher Scientific) together with Probenecid (Thermo Fisher Scientific) in 37 °C for 30 min. After the cells were added with indicated reagents, changes of intracellular calcium were immediately monitored (emission at 515 nm, excitation at 488 nm) by using Synergy H1 multi-mode reader (Bio Tek, Winooski, USA) every 5 s.

Mitochondrial calcium fluorescence was measured by 2.5 µM Rhod-2AM (Thermo Fisher Scientific) together with Pluronic F-127 (Sigma-Aldrich) in 37 °C for 30 min. After the cells were added with indicated reagents, changes of mitochondrial calcium were immediately monitored (emission at 581 nm, excitation at 552 nm) by using Synergy H1 multi-mode reader (Bio Tek) every 5 s.

### Mitochondrial Ca^2+^ uptake

Mitochondria from Jurkat T cells were isolated by using cell mitochondria isolation kit (Beyotime) according to the manufacturer’s instruction. Mitochondria were suspended in experiment buffer containing 1 mM succinate, 2 µM rotenone, 1 µM MgCl_2_, 1 mM ATP and 0.1 µM calcium probe Ca^2+^ Green 5N (Thermo Fisher Scientific) in 37 °C for 15 min. The calcium fluorescence was measured at 506 nm excitation and 532 nm emission by using Synergy H1 multi-mode reader (Bio Tek). 25 µM CaCl_2_ was added every 3 min continuously until Ca^2+^ accumulation capacity was exhausted.

### Real-time PCR

The total RNA of the cell samples was extracted with 1000-800 µl RNAiso Plus reagent (Takara, Kyoto, Japan) according to manufacturer’s instructions, and cDNA was synthesized using the HiScript III RT SuperMix (Vazyme, Nanjing, China). Sequences of qPCR primers were from PrimerBank followed by NCBI blast. mRNA levels were normalized to β-actin mRNA and expressed as fold change relative to the control or vehicle group. Detailed primers sequences shown in **Appendix Table S1**.

### Co-culture of CD8^+^ T cells and HBV transfected Hepa1-6 cells and assessment of HBV replication

The CD8^+^ T cells (effector cells) were added to the confluent Hepa1-6 cells (target cells). The CD8^+^ T cells and Hepa1-6 cells were co-culture for 48 h using E:T ratios of 1:5. At same time, Hepa1-6 cells transfected with the rAAVdj-HBV 1.3 (MOI 1e3). At the end of the experiments, the targets cells and the corresponding culture supernatants were harvested for measurement of Hbx DNA quantitation.

### Western Blotting

The collected cells were lysed in RAPI buffer containing with 1mM PMSF to extract total protein. Nuclear protein and cytosolic protein were extracted from the Nuclear and Cytoplasmic Protein Extraction Kit (Beyotime). The protein concentration was detected by the BCA assay (Beyotime). 60-80 µg proteins were subjected into SDS polyacrylamide gel (Bio-Rad, Hercules, CA, USA) and transferred to PVDF membranes (Bio-rad). Then the membranes were blocked with 5% milk in TBST for 1 h at 37 °C, followed by incubation with primary antibodies and HRP-conjugated secondary antibodies. The membranes bands were visualized with ECL reagent (Bio-rad), the signals were captured by using an iBright CL1000 System (Invitrogen, USA).

### Immunoprecipitation

The collected cells were lysed in NP-40 buffer supplemented with 1mM PMSF and phosphatase inhibitors. Lysates were cleared by Protein A/G Agarose (Thermo Fisher Scientific) for 1 hour and followed by immunoprecipitated with primary antibody of PP2B-Aα, Stim1, Phosphoserine and Phosphothreonine or IgG control antibody for 4-6 h at 4 °C. The mixture was washed with lysis buffer reducing sample buffer, and the proteins were eluted by boiling 15 min in SDS sample buffer then subjected to western blot analysis.

### Elisa

Cytokine concentration in liver homogenates were detected using mouse IFN-γ kits (ExCell Bio, Shanghai, China) according to the manufacturer’s instructions. The concentration of total protein in homogenates was determined by BCA protein assay kit (Beyotime Biotechnology, Nantong, China) to normalize the cytokines concentration. Cytokine concentration in serum was detected using mouse HBsAg Elisa kits (Enzyme-linked Biotechnology, Shanghai,China) according to the manufacturer’s instructions. The absorbance of each sample was then measured at 450 nm using Synergy H1 Microplate Reader (Bio-Tek).

### Serology for Transaminase Activity

Liver function biomarkers of mice were analyzed by Alanine aminotransferase Assay Kit (C009-2-1) and Aspartate aminotransferase Assay Kit (C010-2-1) from Nanjing Jiancheng Bioengineering Institute (Nanjing, China), according to the manufacturer’s instructions. The absorbance of each sample was then measured at 510 nm using Synergy H1 Microplate Reader (Bio-Tek).

### Histology and immunohistochemistry

Tissue specimens were fixed in 10% formalin for 12–24 h, dehydrated and paraffin embedded. Hematoxylin-eosin (HE) staining of mice liver sections (5 µm) was used to evaluate mice liver inflammation.

Standard immunohistochemical procedures were performed. Tissue sections were incubated with the primary rabbit antibody to HBsAg (1:100; MXB Biotechnologies, Fuzhou, China) for 2 h at room temperature, followed by 30 min incubation with the peroxidase-conjugated secondary antibody (1:400). All images were acquired using Leica system (DM6B, Heidelberg, Germany).

### Statistical analysis

Data are presented as Mean ± SEM; number of replicates is stated in the figure legends. Three independent experiments were performed throughout in this study. Animal studies were age and sex matched and randomized by body weight. Significance between 2 groups were analyzed with student’s *t*-test. Representative graphs were used GraphPad Prism 8 software. Flow cytometry was carried out using BD Celesta and analyzed with FlowJo software V10. Metabolic analyses were used MetaboAnalyst 3.0. P values of < 0.05 were considered significant. ***p<0.001, **p < 0.01, and *p < 0.05.

## Acknowledgments

This work was financially supported by National Natural Science Foundation of China (grants 81720108032, 81930109, 81973559 and 82173886); the National Key Research and Development Programme of China (2021YFA1301300); Overseas Expertise Introduction Project for Discipline Innovation (G20582017001) and Sanming Project of Medicine in Shenzhen (grants SZSM201801060 to H. Hao).

## Author contributions

Haiping Hao and Lijuan Cao designed the study; Chujie Ding, Yu Hong and Yuan Che performed the most experiments and collected and analyzed the data, Chujie Ding, Yu Hong and Yuan Che contributed equally to this manuscript; Haiping Hao, Lijuan Cao and Chujie Ding wrote and revised the manuscript.

## Conflicts of interest

The authors declare no conflicts of interest.

## Expanded View Figures and Legends

**Figure EV1.**
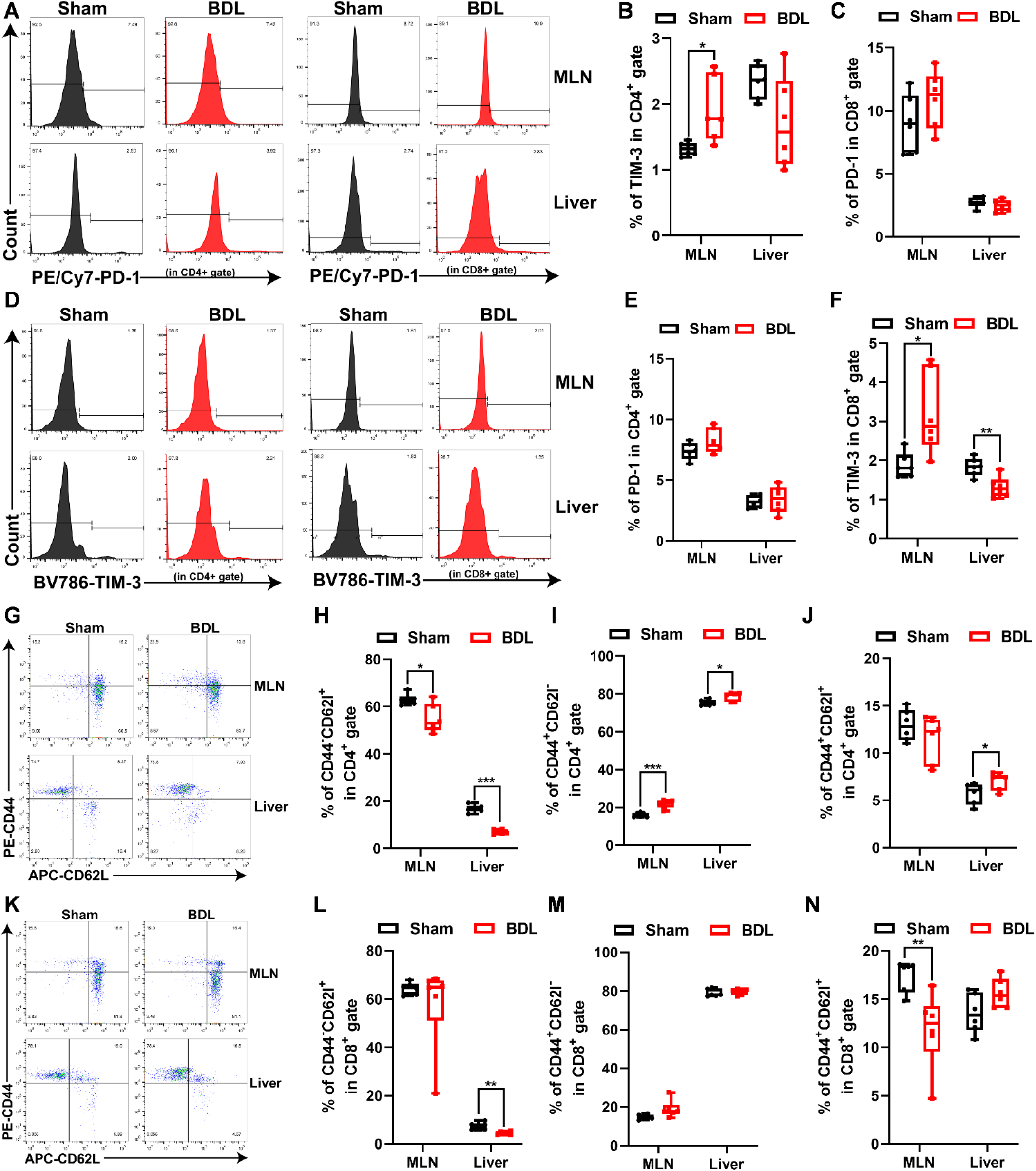
The effect of cholestasis on T cells exhaustion and differentiation. Mice were performed with BDL surgery for 5 days before lymphocytes in MLN and liver were analyzed by flow cytometry. A-C Flow cytometry analysis (A) and quantitative data of PD-1 among CD4^+^ T cells (B) and CD8^+^ T cells (C) in lymphocytes isolated from MLN and liver (*n* = 6). D-F Flow cytometry analysis (D) and quantitative data of TIM-3 among CD4^+^ T cells (E) and CD8^+^ T cells (F) in lymphocytes isolated from MLN and liver (*n* = 6). G-J Flow cytometry analysis (G) and quantitative data of CD4^+^ naive T cells (H, CD44^-^CD62l^+^), effector T cells (I, CD44^+^CD62l^-^) and memory T cells (J, CD44^+^CD62l^+^) in lymphocytes isolated from MLN and liver (*n* = 6). K-N Flow cytometry analysis (K) and quantitative data of CD8^+^ naive T cells (L, CD44^-^CD62l^+^), effector T cells (M, CD44^+^CD62l^-^) and memory T cells (N, CD44^+^CD62l^+^) in lymphocytes isolated from MLN and liver (*n* = 6). Data information: Data are representative of three independent experiments and expressed as mean ± SEM. ***p<0.001, **p < 0.01, *p < 0.05 using two-tailed Student’s *t*-test.

**Figure EV2.**
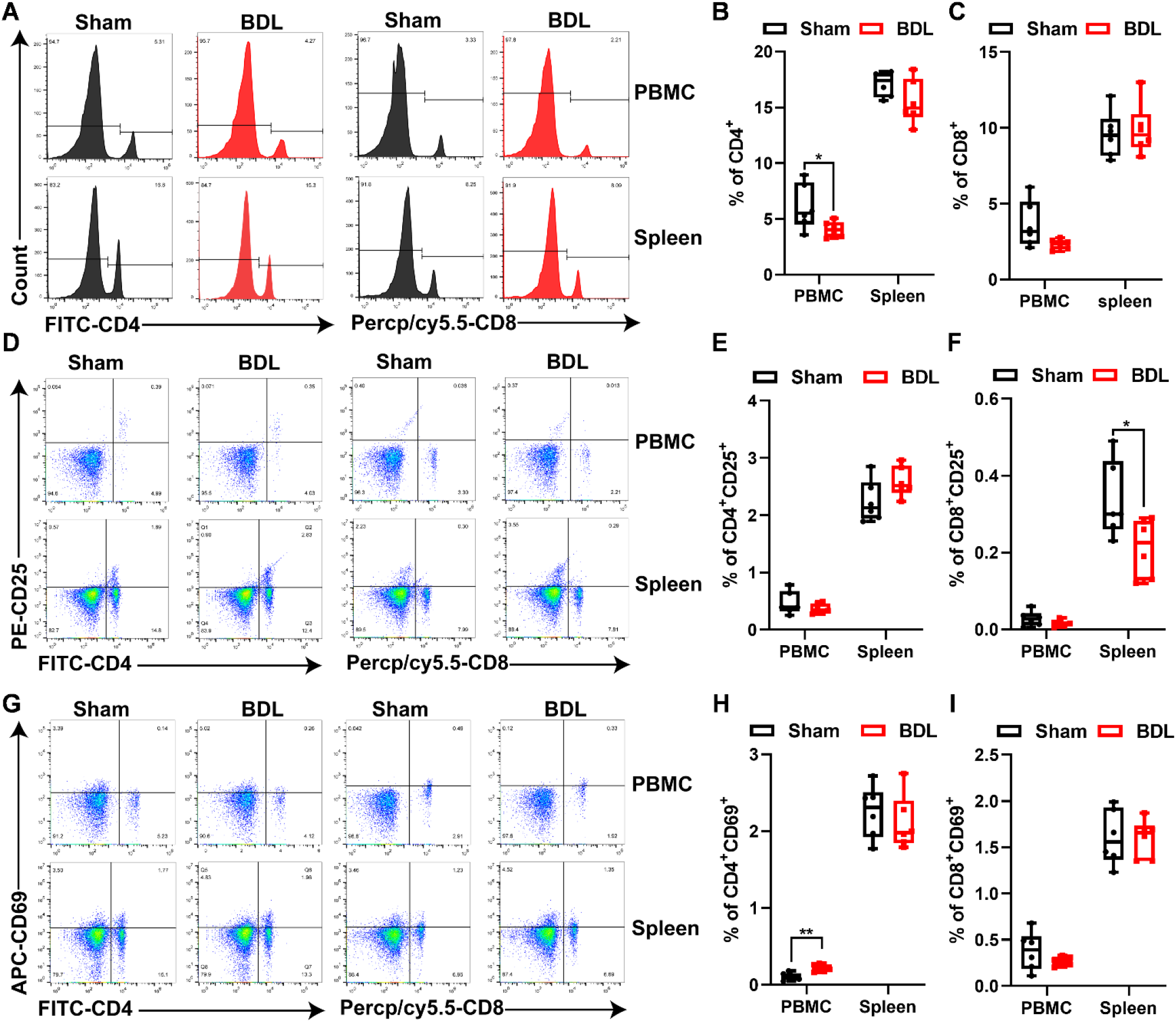
Cholestasis does not affect number and activation in PBMC and spleen. Mice were performed with BDL surgery for 5 days before lymphocytes in PBMC and spleen were analyzed by flow cytometry. A-C Flow cytometry analysis (A) and quantitative data of CD4^+^ (B) and CD8^+^ (C) T cells in PBMC and spleen (*n* = 6). D-F Flow cytometry analysis (D) and quantitative data of CD4^+^CD25^+^ (E) and CD8^+^CD25^+^ (F) in PBMC and spleen (*n* = 6). G-I Flow cytometry analysis (G) and quantitative data of CD4^+^CD69^+^ (H) and CD8^+^CD69^+^ (I) in PBMC and spleen (*n* = 6). Data information: Data are representative of three independent experiments and expressed as mean ± SEM. ***p<0.001, **p < 0.01, *p < 0.05 using two-tailed Student’s *t*-test.

**Figure EV3.**
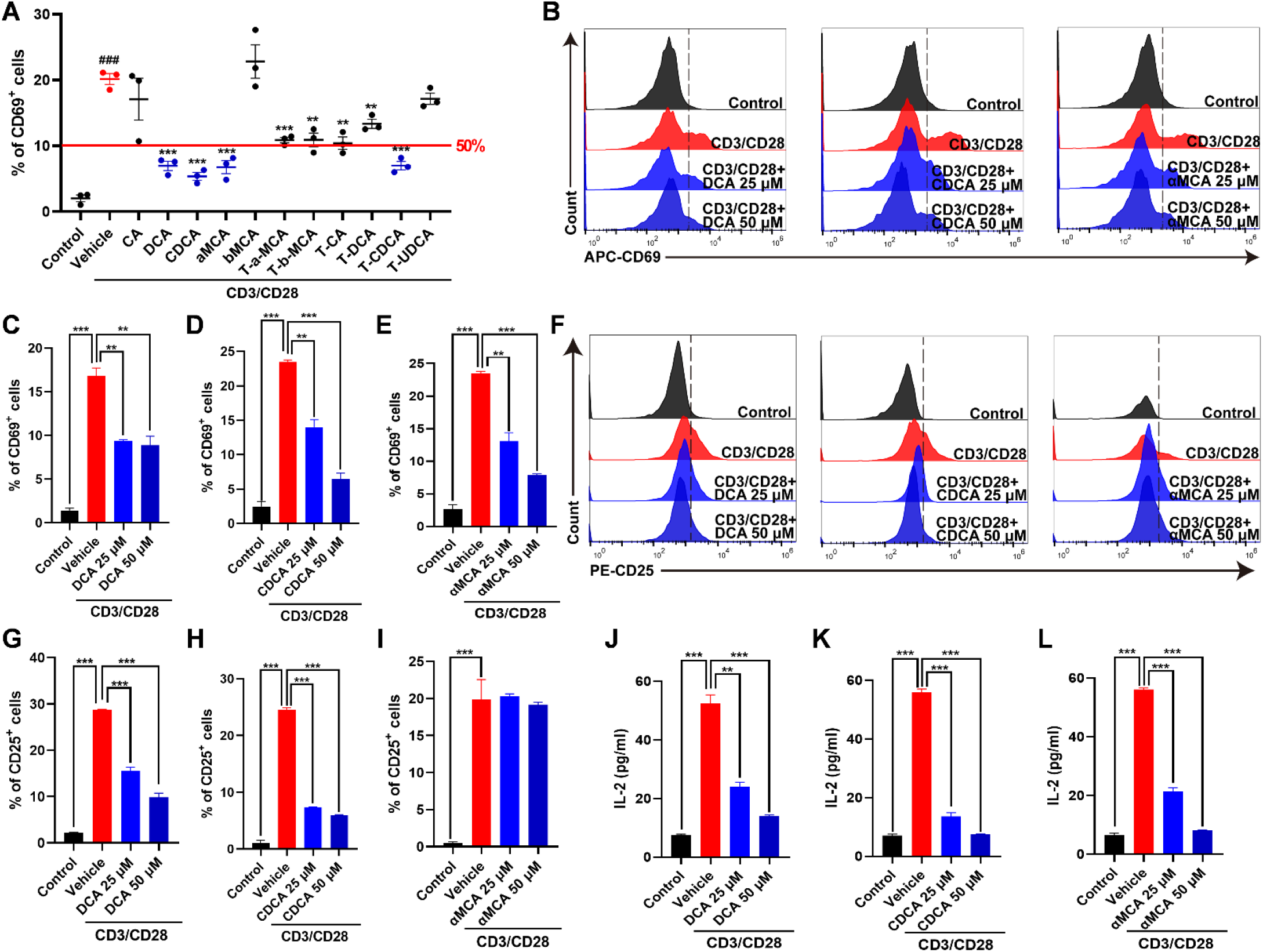
Bile acids control mouse primary CD4^+^ T cells activation *in vitro*. A Primary mouse CD4^+^ T cells were stimulated with CD3/CD28 for 6 h in the presence of various species of bile acids at 50 µM concentration and proportion of CD69^+^ cells was detected by flow cytometry. B-E Primary mouse CD4^+^ T cells were stimulated with CD3/CD28 for 6 h in the presence of DCA, CDCA and α-MCA at 25 or 50 µM. Flow cytometry analysis (B) and quantitative data of CD69^+^ cells were shown in (C) for DCA, (D) for CDCA, (E) for α-MCA. F-I Primary mouse CD4^+^ T cells were stimulated with CD3/CD28 for 24 h in the presence of DCA, CDCA and α-MCA at 25 or 50 µM. Flow cytometry analysis (B) and quantitative data of CD25^+^ cells were shown in (G) for DCA, (H) for CDCA, (I) for α-MCA. J-L ELISA analysis of supernatant IL-2 concentrations in primary mouse CD4^+^ T cells. Primary mouse CD4^+^ T cells were stimulated with CD3/CD28 for 24 h in the presence of DCA (J), CDCA (K) and α-MCA (L) at 25 or 50 µM. Data information: Data are representative of three independent experiments and expressed as mean ± SEM (*n* = 3). (A) In comparison with the control group, ^###^p<0.001; compared with the vehicle group ***p<0.001, **p < 0.01, *p < 0.05 between the indicated groups; two-tailed Student’s *t*-test. (B-L) ***p<0.001, **p < 0.01, *p < 0.05 between the indicated groups using two-tailed Student’s *t*-test. **See also Figure EV4.**

**Figure EV4.**
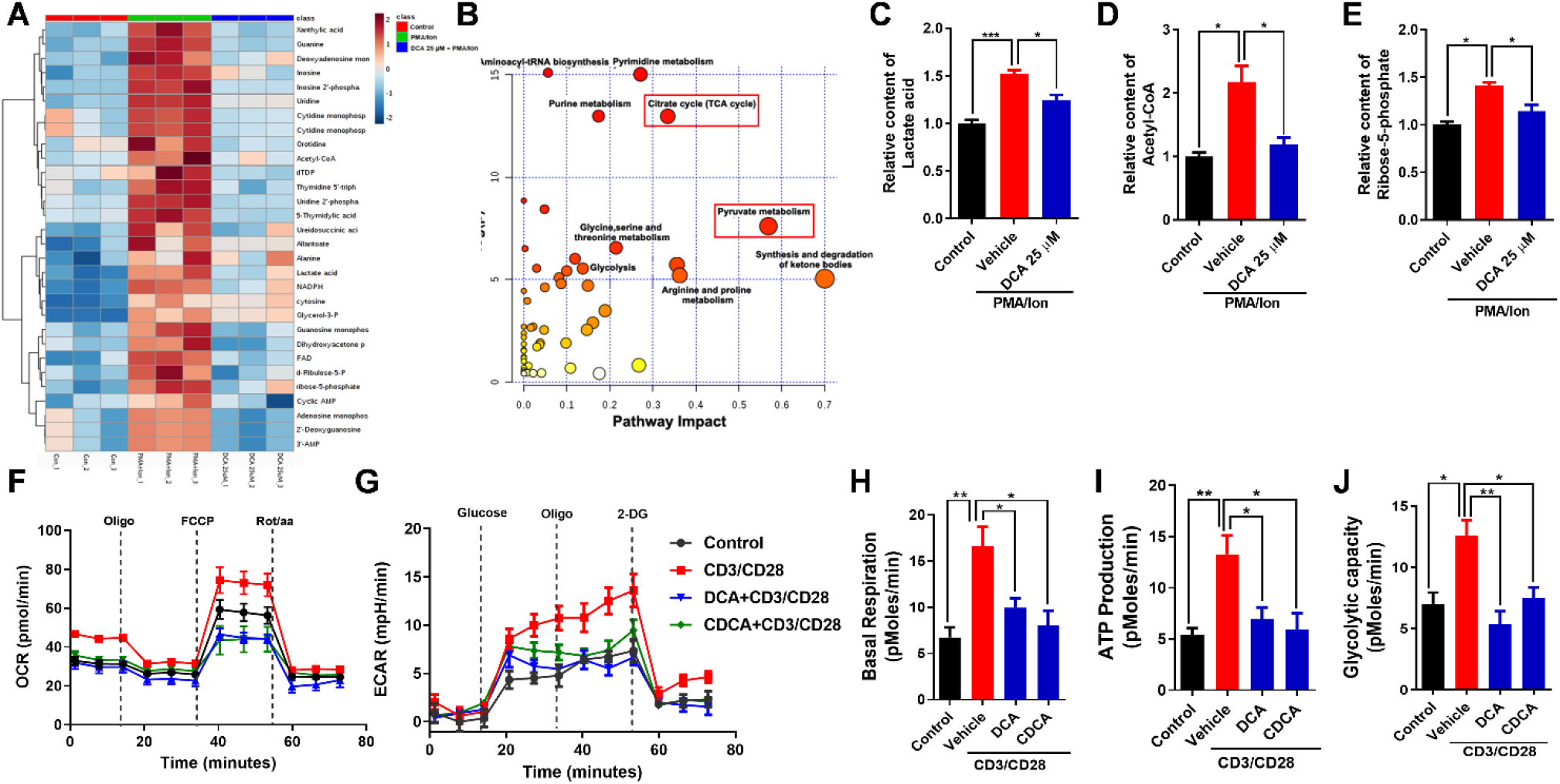
Bile acids regulate metabolism reprogramming of T cells. A Analysis of polar metabolites by LC-MS/MS in Jurkat cells. Jurkat cells were pretreated with DCA at 25 µM for 24 h and then stimulated with PMA/Ion for 6 h. A heatmap shows differences of identified potential biomarkers between the DCA-treated cells and the control group. B Metabolic pathway enrichment analysis of potential biomarkers identified from the DCA-treated cells. C-E Representative metabolites lactate acid (C), acetyl-CoA (D) and ribose-5-phosphate (E) that underwent significant level changes were summarized respectively. F-J Real-time changes in the ECAR (F), OCR (G), basal respiration (H), ATP production (I) and glycolytic capacity (J) of primary CD4^+^ T cells stimulated with anti-CD3/anti-CD28 for 24 h in the presence of 25 µM DCA or CDCA. Data information: Data are representative of three independent experiments and expressed as mean ± SEM. ***p<0.001, **p < 0.01, *p < 0.05 using two-tailed Student’s *t*-test.

**Figure EV5.**
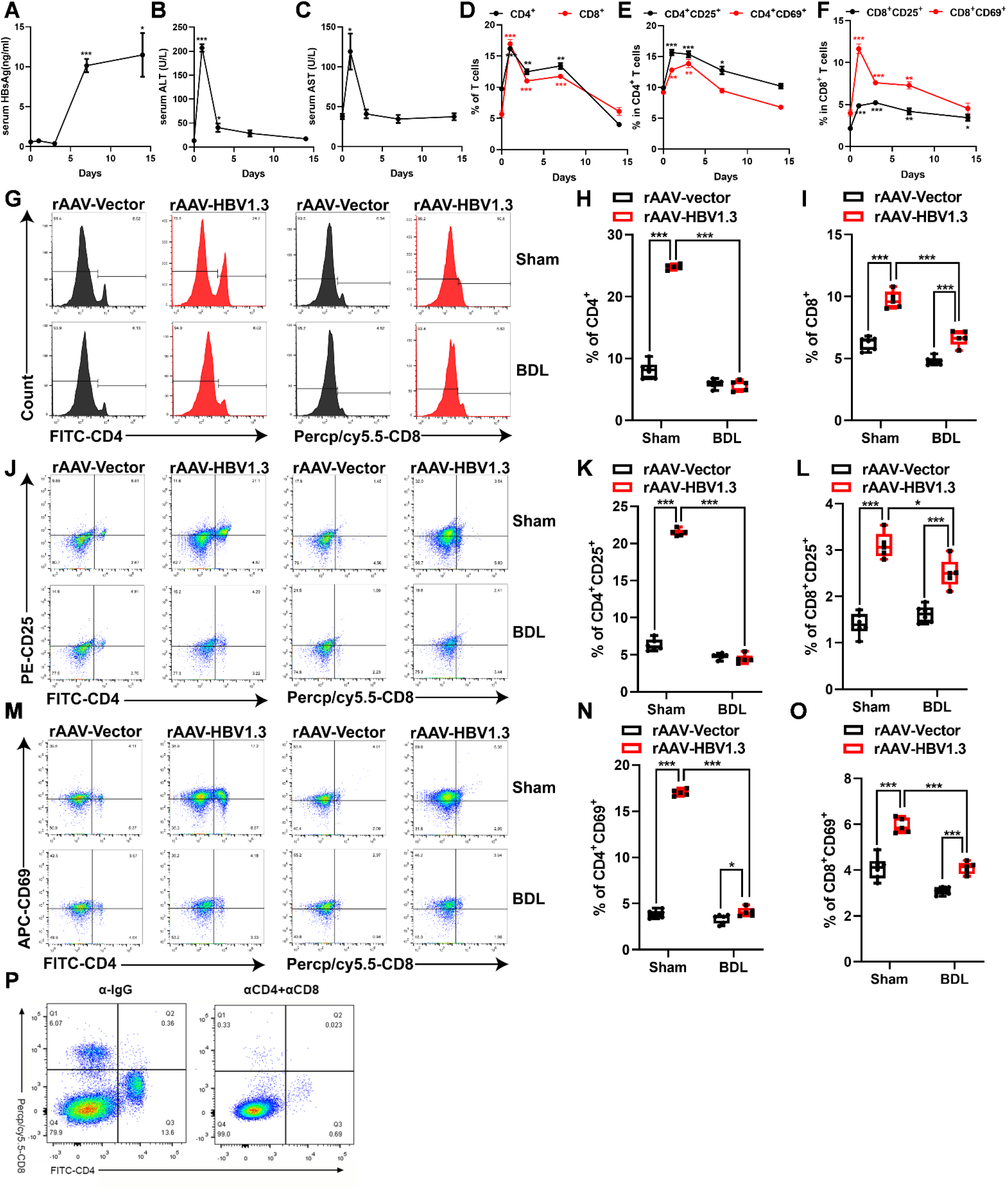
Anti-viral T cell immunity is defective in cholestasis mice. A-C Mice were hydrodynamically transfected with rAAV-HBV1.3 or rAAV-Vector. Serum HBsAg (A), ALT (B) and AST (C) levels in HBV transfected mice over time was detected (*n* = 3). D-F The percentage of CD4^+^ and CD8^+^ T cells (D) and proportion of CD25^+^ and CD69^+^ in CD4^+^ T cells (E) and CD25^+^ and CD69^+^ in CD8^+^ T (F) cells from HBV transfected mice liver lymphocytes were detected at the indicated time points (*n* = 3). G-I Representative flow cytometry analysis (G) and quantitative data of CD4^+^ (H) and CD8^+^ (I) T cells in liver lymphocytes at 1^st^ day after rAAV-HBV1.3 virus transfection with BDL mice (rAAV-Vector, *n* = 6; rAAV-HBV1.3, *n* = 5). J-L Representative flow cytometry analysis (J) and quantitative data of CD4^+^CD25^+^ (K) and CD8^+^CD25^+^ (L) in liver lymphocytes at 1^st^ day after rAAV-HBV1.3 virus transfection with BDL mice (rAAV-Vector, *n* = 6; rAAV-HBV1.3, *n* = 5). M-O Representative flow cytometry analysis (M) and quantitative data of CD4^+^CD69^+^ (N) and CD8^+^CD69^+^ (O) in liver lymphocytes at 1^st^ day after rAAV-HBV1.3 virus transfection with BDL mice (rAAV-Vector, *n* = 6; rAAV-HBV1.3, *n* = 5). P Representative flow cytometry plots, showing relative abundance of CD4^+^ and CD8^+^ T cells in PBMC at 7^th^ day after rAAV-HBV1.3 virus transfection with treatment of αIgG or αCD4+αCD8 depleting antibodies in combination. Data information: Data are representative of three independent experiments and expressed as mean ± SEM. ***p<0.001, **p < 0.01, *p < 0.05 using two-tailed Student’s *t*-test.

